# Collapse and rescue of evolutionary food webs under global warming

**DOI:** 10.1101/701839

**Authors:** Youssef Yacine, Korinna T. Allhoff, Avril Weinbach, Nicolas Loeuille

**Affiliations:** Institute of Ecology and Environmental Sciences Paris (iEES Paris), Sorbonne Université/CNRS/IRD/INRA/Université de Paris/UPEC, 4 place Jussieu, 75252 Paris Cedex 5, France; University of Tübingen, Institute of Evolution and Ecology, Auf der Morgenstelle 5, D-72076, Tübingen, Germany

**Keywords:** Adaptive Dynamics, Biodiversity conservation, Body mass evolution, Climate Change, Co-evolution, Consumer-Resource model, Eco-evolutionary tipping point, Evolutionary rescue, Trophic networks

## Abstract

1. Global warming is severely impacting ecosystems and threatening ecosystem services as well as human well-being. While some species face extinction risk, several studies suggest the possibility that fast evolution may allow species to adapt and survive in spite of environmental changes.
2. We assess how such evolutionary rescue extends to multitrophic communities and whether evolution systematically preserves biodiversity under global warming.
3. More precisely, we expose simulated trophic networks of co-evolving consumers to warming under different evolutionary scenarios, which allows us to assess the effect of evolution on diversity maintenance. We also investigate how the evolution of body mass and feeding preference affects coexistence within a simplified consumer-resource module.
4. Our simulations predict that the long-term diversity loss triggered by warming is considerably higher in scenarios where evolution is slowed down or switched off completely, indicating that eco-evolutionary feedback indeed helps to preserve biodiversity. However, even with fast evolution, food webs still experience vast disruptions in their structure and functioning. Reversing warming may thus not be sufficient to restore previous structures.
5. Our findings highlight how the interaction between evolutionary rescue and changes in trophic structures constrains ecosystem responses to warming with important implications for conservation and management policies.

## Introduction

The consequences of global change on biodiversity are now well-documented. They include the extinction of species (Barnosky et al., 2011), changes in species demography, ranges and phenologies (Parmesan & Yohe, 2003) and alterations of ecological interactions (Tylianakis et al., 2008). Significant ecological network reorganizations and changes in ecosystems functioning are therefore expected. In such a stressful context, it is unclear whether the evolution (or co-evolution) of species will have a net positive or negative effect on network maintenance and stability.

Existing models investigating ecosystem responses to global warming often ignore evolutionary processes and the interaction of species. This is particularly true for “niche” or “envelope” models (Colwell & Rangel, 2009) that link data of species occurrence with climatic variables to build a multivariate statistical representation of the species niche. Future distributions of species are then predicted according to different climate change scenarios (e.g., Pearson et al., 2002) implicitly assuming that niches are fixed. Evolution, however, affects species’ fundamental niches while the reshuffling of species interactions can lead to changes in realized niches (Tylianakis et al., 2008). While envelope models are important first steps, they are thus limited in their ability to understand the effects of global warming, and to provide relevant conservation policies (Lavergne et al., 2010).

Ignoring evolution is based on the controversial assumption that ecological and evolutionary processes act on separate timescales. Recent studies, however, indicate that evolution may act within a few generations (Koch et al., 2014; Olsen et al., 2004), especially in a context of anthropogenic pressures and disturbances (Hendry, Farrugia, & Kinnison, 2008). For example, many empirical studies (e.g., Daufresne, Lengfellner, & Sommer, 2009) document warming-induced reductions in body size (reviewed in Sheridan & Bickford, 2011). While a clear identification of evolution vs other processes of phenotypic variations is often lacking, these empirical variations suggest that global warming exerts strong selective pressures on body size. This is crucial because body size is a key biological trait influencing many ecological constraints (Brown et al., 2004), including trophic and competitive interactions. Therefore variations of body size are likely to affect the whole food web (e.g., O’Gorman et al., 2017). Next to body size, changes in foraging strategies have also been documented in the context of global change, potentially resulting in a rewiring of the corresponding community network. Several studies, for instance, documented contemporary evolution in the diet of native herbivores to include invasive plant species (e.g., Carroll et al., 2005). Another example is the butterfly *Aricia agestis* (UK), which widened its larval host range as a result of increased temperatures (Pateman et al., 2012). The associated genetic signature (Buckley, Butlin, & Bridle, 2012) strongly suggests this is a case of evolutionary adaptation, that has enabled the species to expand its range poleward.

In some situations, evolution enables species to survive environmental deterioration. Gomulkiewicz & Holt (1995) theorized this concept as “evolutionary rescue”: adaptive evolutionary change restores positive growth in declining populations and prevents extinction. The empirical evidence for evolutionary rescue, as well as the different factors involved, are discussed in detail in the review by Carlson, Cunningham, & Westley (2014). Effects of evolution are however not always positive for biodiversity. Environmental change can alter evolutionary dynamics so that a non-viable phenotype is selected given the new ecological conditions (“evolutionary trapping”, Ferriere & Legendre, 2012; empirical e.g., Singer & Parmesan, 2018). Negative effects of evolution on diversity also arise when the evolution of one species drives its interacting partners towards extinction (“evolutionary murder”, e.g., Dieckmann, Marrow, & Law, 1995). Assessing both positive and negative effects of evolution is crucial to implement relevant conservation decisions (Stockwell, Hendry, & Kinnison, 2003).

The last example readily highlights how accounting for the community context is equally important, since the interplay between evolution and species interactions can lead to unexpected behaviors, as illustrated by the two following examples. Within a spatially explicit model of competitive communities, Norberg et al. (2012) demonstrated that evolution under climate change can create extinction debts long after climate stabilization, but only when competition is accounted for. Osmond, Otto, & Klausmeier (2017) showed that, surprisingly, the maximum environmental change rate an evolving prey population can support increases in the presence of predators. In some cases, the presence of a predator accelerates prey evolution, facilitating its persistence (e.g., selective predation on maladapted prey).

Within a network context, the conditions for and the mechanisms underlying evolutionary rescue can therefore be much more complex compared to monospecific systems. A focal species could be rescued by the evolution of another interacting species (“indirect evolutionary rescue”, Yamamichi & Miner, 2015). On the other hand, if evolutionary rescue happens for various species within the network, a focal species could still go extinct if its enemies’ (resp. positive interactors) recovery is too fast (resp. too slow) compared to its own (Loeuille, 2019). A key question is therefore whether evolutionary rescue, derived from a monospecific approach, can extend to ecological networks. Single or few species models represent important stepping stones to address this question, because they focus on essential key mechanisms. However, they provide limited insights into the complex indirect interactions occurring in diverse networks (Ellner & Becks, 2011). Although currently rare (but see Norberg et al. (2012)), models that consider evolutionary processes within multispecies communities are thus essential.

Here, we ask whether evolutionary rescue at the network scale can promote food web persistence under warming. We address this question using an evolutionary food web model that is based on body masses and feeding preferences. Starting from a single consumer population feeding on a basal resource, the evolution of these two consumer traits leads to the emergence of complex multitrophic communities. Related models have been shown to share many features with empirical food webs (Allhoff et al., 2015; Loeuille & Loreau, 2005). Once the initial build-up is complete, we expose such simulated networks to warming events. We vary both the warming intensity and the mutation rate in order to investigate the interplay between warming, evolution and network diversity maintenance. While this simulation approach provides valuable insights into eco-evolutionary responses to warming at the community scale, it is admittedly difficult to unravel the underlying mechanisms governing these responses. We therefore complement our work by analytically investigating the processes by which evolution facilitates or constrains diversity maintenance within a simplified framework consisting of only one consumer and its resource.

The overarching goal of the present study is to expand existing knowledge from simplified mono-specific systems to complex multispecies networks. We show that with or without evolution, warming is responsible for considerable diversity losses. In line with both theoretical (Binzer et al., 2012; Weinbach et al., 2017) and experimental work (Petchey et al., 1999), we find these losses to be more frequent among upper trophic levels. Evolution has a positive effect on diversity maintenance. It notably enables diversity to progressively and partially recover after a transient collapse. Our results are globally in line with the expectations derived from evolutionary rescue theory (Gomulkiewicz & Holt, 1995).

## Model

### Ecological dynamics

The trophic network consists of a set of primary producers providing energy to the whole community (hereafter basal resource of (aggregated) biomass density *B*_0_) and consumer morphs *i*. Its dynamics follow:

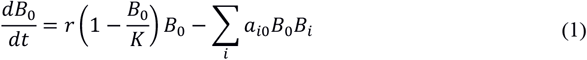

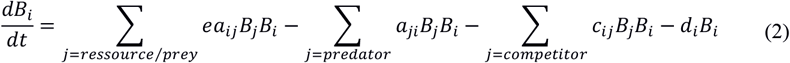

The basal resource follows a logistic growth in the absence of higher trophic levels, with intrinsic growth rate *r* and carrying capacity *K* (equation 1). Variations of biomass density *B_i_* of a morph *i* (equation (2)) are composed of four terms: predation gain, predation loss, losses due to interference competition, as well as intrinsic losses due to respiration and basic mortality (hereafter respiration). A complete list of parameters and variables is provided in Table 1.

**Table 1:**
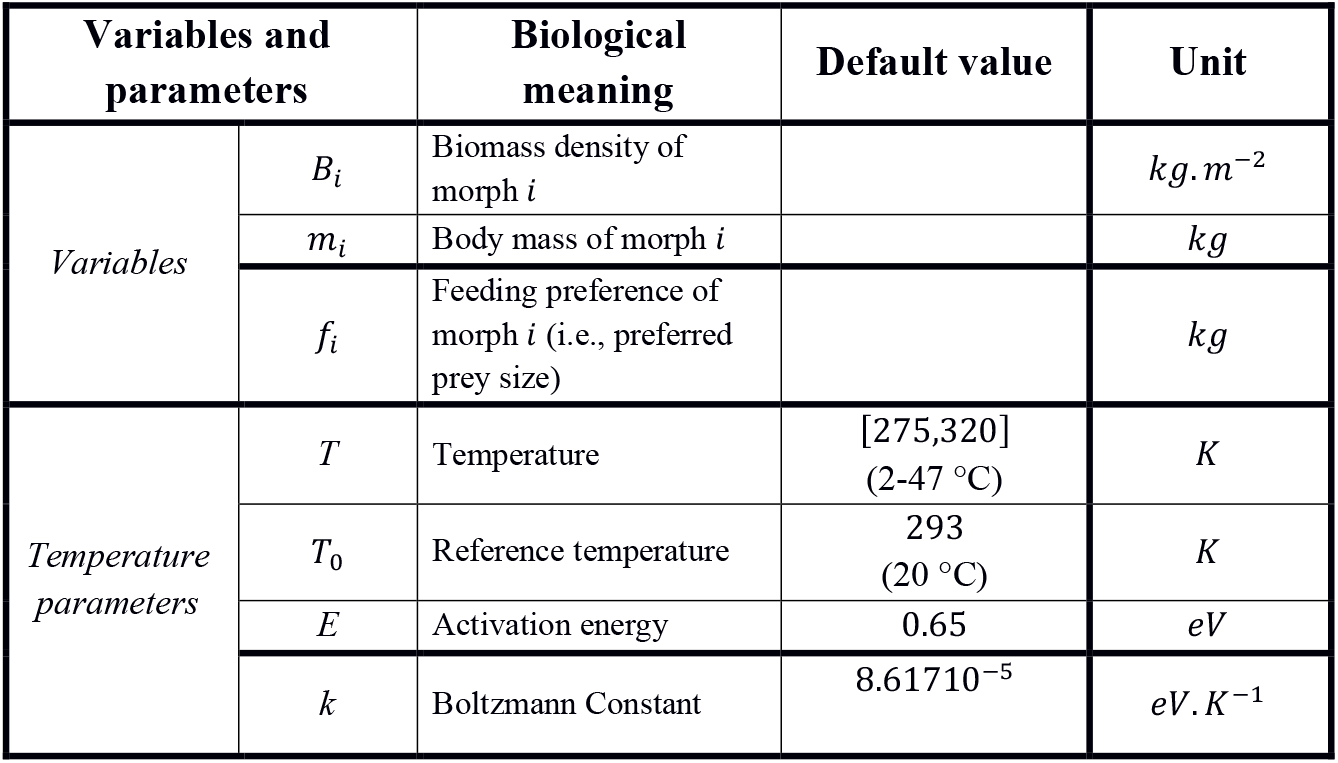
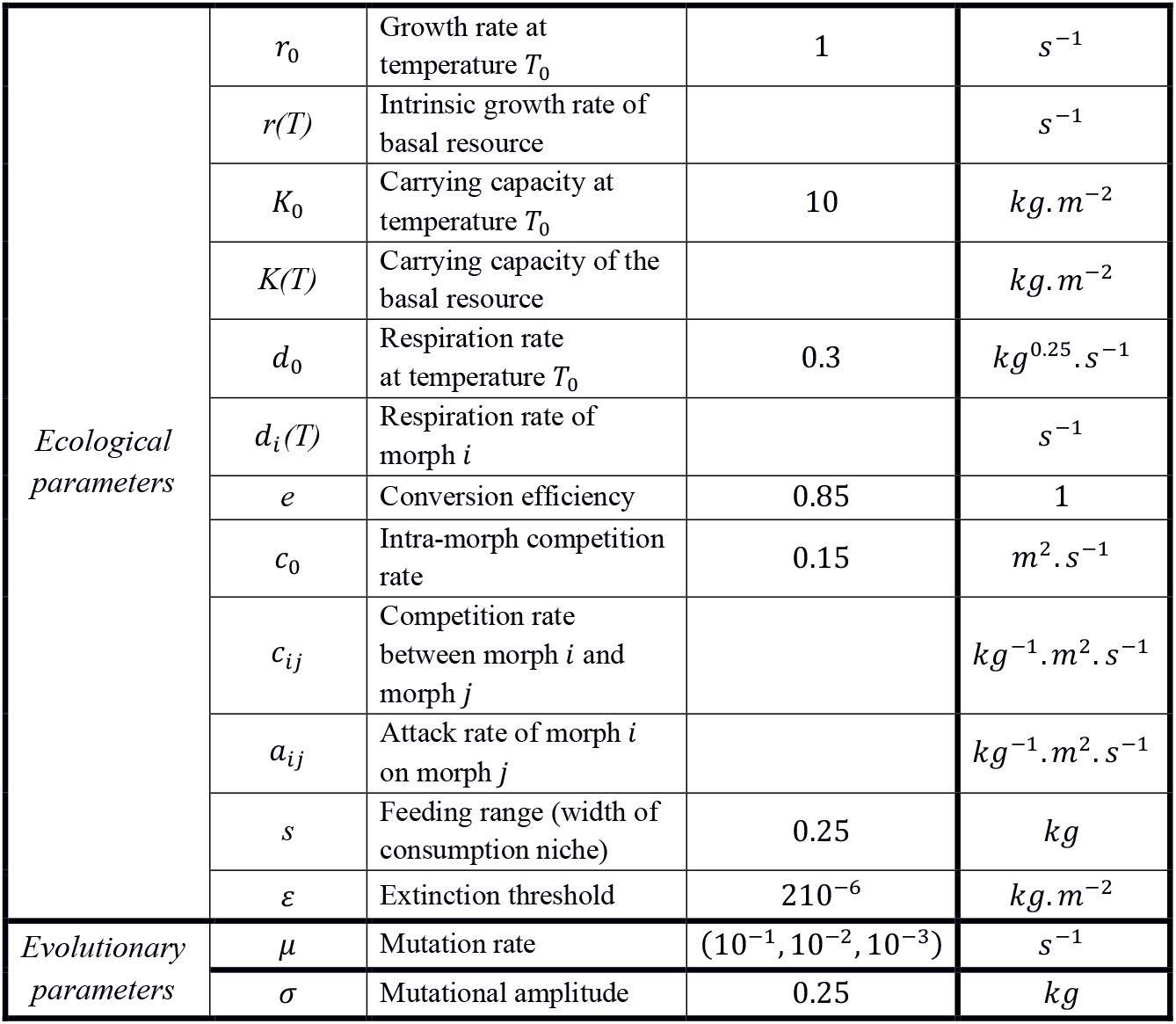
Variables and parameters. Dependence on temperature is explicitly indicated. The values given here represent the standard parameter set used in our simulations, unless stated otherwise.

### Temperature dependence

Biochemical reactions and thus metabolic rates are known to grow exponentially with temperature. We therefore incorporate temperature dependency as an Arrhenius function in the resource growth rate *r* and the respiration rate *d_i_* of consumer *i* following Brown et al. (2004):

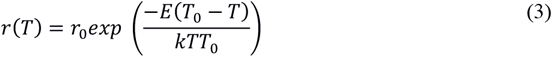

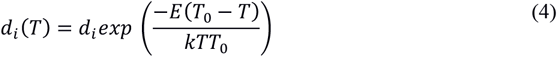

We furthermore assume that the carrying capacity of the basal resource decreases exponentially with temperature as in Fussmann et al. (2014).

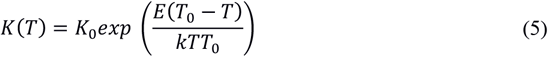

Here, *k* is the Boltzmann constant, *E* the activation energy and *T*_0_ the reference temperature (Table 1). We assume these relationships to hold over the temperature range we consider here (275-300 K, 2-47 °C) (but see Discussion).

### Presentation of evolving traits

Each consumer morph *i* has two adaptive traits: body mass *m_i_* and feeding preference *f_i_*. Body mass is known to largely constrain trophic interactions (see for instance Woodward et al. (2005)). The basal resource has a body mass *m*_0_ that we assume fixed. The feeding preference *f_i_* corresponds to the prey body mass allowing a maximal attack rate. Fig. 1 illustrates how traits constrain trophic interactions.

**Figure 1:**
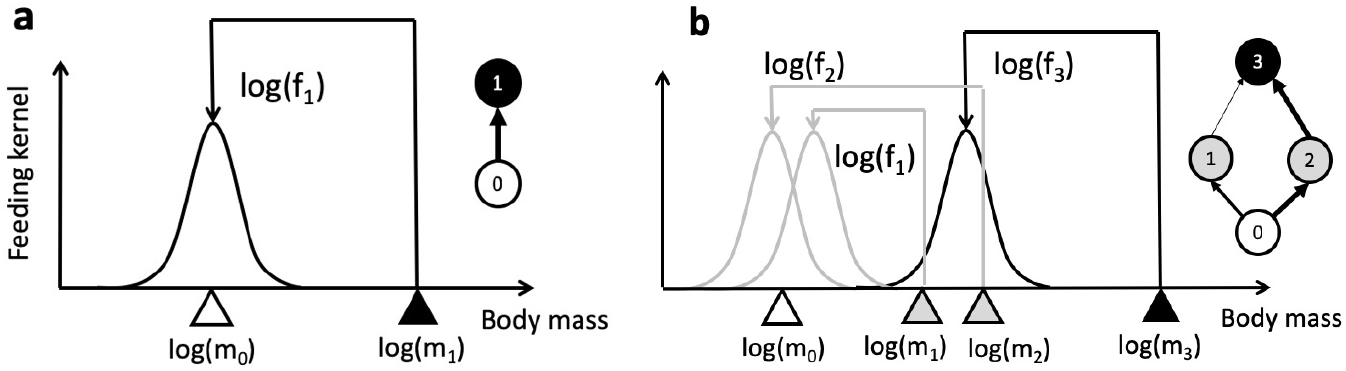
(Adapted from Allhoff et al., 2015). **a.** Consumer-Resource module. The consumer 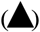 has a body mass ***m*_1_** and feeds on the basal resource 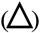 with maximum attack rate (black curve) because its feeding preference ***f*_1_** corresponds to the resource body mass **m_0_**. Resulting trophic interaction on the right. **b.** Complex multitrophic network emerging by co-evolution. The snapshot here shows 3 morphs: morph 3 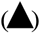 feeds on morph 1 and 2 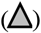 with respectively low and high attack rates, morph 1 and 2 feed on the basal resource 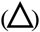 with respectively low and high attack rates. Resulting trophic module on the right. This is a snapshot view; the real networks are dynamic and typically have many more morphs.

Respiration and attack rates scale with body mass (Brown et al., 2004; Peters, 1983). In our model, attack rates depend on the relative differences between predators’ feeding preferences and prey’s body masses. Supporting our formulation, several empirical studies indicate that predation intensity is determined by predator-prey body mass ratios (Naisbit et al., 2011; Vucic-Pestic et al., 2010). Around a predator’s feeding preference, attack rates are thus distributed right-skewed on an absolute scale, meaning higher predation on smaller prey (Aljetlawi, Sparrevik, & Leonardsson, 2004; Brose et al., 2008). We furthermore assume competition to increase for similar feeding preferences, due to niche overlap (Macarthur & Levins, 1967). The feeding niche width is denoted by *s* (Fig. 1) and *c*_0_ is a scaling constant for the interference competition.

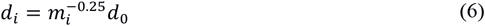

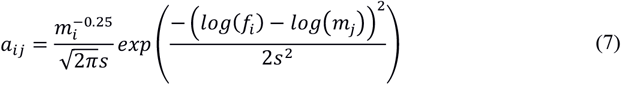

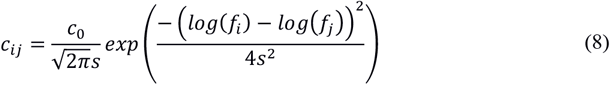

## Methods

### Simulations

Simulations start from a single ancestor morph (*m*_1_ = 100, *f*_1_ = 1) feeding on the basal resource (*m*_0_ = 1). The succession of mutation events (one mutation every 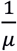 time steps) leads to the emergence of a multitrophic network with typically 30 to 40 morphs and 4 to 5 trophic levels (Allhoff et al., 2015; Loeuille & Loreau, 2005). Proportionally to population abundances, a parent morph is chosen randomly at each mutation event. Mutant traits are then drawn from log-normal distributions centered around the parent’s traits. Occasional big mutational steps are allowed (details in Appendix S2.B). The mutant is initially very rare as its initial biomass density corresponds to the extinction threshold *ε* taken from the parent population.

To assess the role of evolution for the maintenance of diversity, we consider two evolutionary scenarios: with evolution (“scenario E”) and without evolution (“scenario NE”). Scenarios NE consist of the following sequence of events: (1) the network is built up with a mutation rate *μ*; (2) evolution stops (*μ* = 0) when a quasi-equilibrium is reached; (3) warming occurs; (4) simulation stops when a new quasi-equilibrium is reached. Scenarios E follow the same sequence except (2). We consider quasi-equilibrium situations to be reached when the relative trait diversity variability over time is below a critical value (see Appendix S2.B). We are thus not focusing on the transient state following perturbation but on the long-term response to warming, when a steady state is reached.

We ran a total of 420 simulations. For each of the two scenarios (with evolution E, without evolution NE), we ran simulations with all factorial combinations of *μ* = 10^−1^, 10^−2^, 10^−3^ and Δ*T* = 0, 2, 4, 6, 8, 10, 20 K or °C (initial temperature is always 275K (2°C)). We ran 10 replicates for each combination of parameters. We focus on the role of *μ* and Δ*T*, because a well-supported expectation from evolutionary rescue theory is that evolution is less likely to save the species when the disturbance is high, or when the genetic variability (here brought by mutation) is low (Gomulkiewicz & Holt 1995, Carlson et al. 2014). We want to test whether these assumptions remain valid in a multispecies context.

### Evolutionary rescue at the multitrophic network scale

Ferriere & Legendre (2012) propose a broad definition of evolutionary rescue that we adapt to our network context: “evolutionary rescue occurs when a population subject to environmental change ‘performs better’ under the operation of evolutionary processes than without these processes”. We measure the network performance as the diversity maintained after warming relative to the diversity that was present before (hereafter “persistence”, see appendix S2.B). Diversity is measured either as trait or species diversity. Trait diversity corresponds to the total number of morphs present at a given time. While this is certainly a valuable measure from a functional and structural point of view, a large focus exists in conservation biology on the preservation of species diversity. Because our model ignores genetic details and focuses on phenotypes, the definition of species is notoriously tricky. For lack of better criteria, we define species as clusters in the phenotypic space (Appendix S2.A). We used statistical models implemented in R-software to compare persistence across scenarios. For each mutation rate, we fitted two ANCOVAs to link trait or species diversity persistence with evolutionary scenarios (evolution E vs no evolution NE) and warming intensities (details in Appendix S2.C). The results of the statistical analysis are summarized in Table S2.

### Identifying mechanisms in a simplified model

The Consumer-Resource (CR) module is derived from equations (1) and (2), assuming one consumer feeding on the resource and potentially its conspecifics (cannibalism).

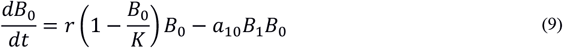

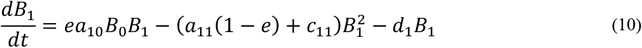

We use adaptive dynamics (Dieckmann & Law, 1996; Metz, Nisbet, & Geritz, 1992) to investigate how warming affects the eco-evolutionary dynamics. Two major assumptions are made: firstly, evolution occurs on longer timescales than ecology and, secondly, mutations are of small amplitude. Timescale separation allows the environment felt by the mutant to be clearly defined by the resident population at ecological equilibrium 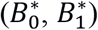 (analytical expression in Appendix S1.A. 1). When the analytical work indicates the system is supposed to go extinct, we assess the potential for evolutionary rescue as in Ferrière & Legendre (2012). More precisely, we undertake simulations to test whether fast evolution may prevent extinction.

For instance, considering trait *m*_1_ (body mass), the invasion fitness 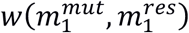 of a mutant corresponds to its relative growth rate in the resident population when rare. Positive values of *w* indicate that the mutant frequency increases, eventually replacing the resident. Evolutionary dynamics are captured by the sequence of trait substitutions, and can be approximated by the canonical equation of adaptive dynamics (Dieckmann & Law 1996). This equation (Appendix S1 B.1) indicates that the trait evolves through time proportionally to the selection gradient, a quantity that captures how the relative fitness 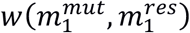 varies with the mutant trait 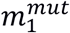 (Appendix S1 B.1). The values of the resident trait 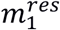 where the selection gradient vanishes are evolutionary singularities. They are classified into Continuously Stable Strategies (CSS, Eshel, 1983), Branching Points (BP, Geritz et al., 1997) and Repellors according to two properties: convergence and invasibility.

Convergence indicates that the trait evolves toward the singularity in its vicinity (convergent: CSS, BP; non-convergent: Repellor). Invasibility specifies whether the singularity may be invaded by nearby mutants (non-invasible: CSS; invasible: BP, Repellor). Branching points are particularly important in terms of diversity: they yield the emergence of a stable dimorphism in the population due to disruptive selection (e.g., the coexistence of two consumer populations with different body masses). Knowing the evolutionary singularities and their properties (Appendix S1 B.3.d) enables the full determination of the evolutionary dynamics of the consumer population within the CR module.

All in all, this simplified framework allows the eco-evolutionary dynamics of the consumer population confronted with warming to be more easily tractable. Such a thorough understanding allows to highlight key mechanisms that may also act at the co-evolving multitrophic scale. However, we keep in mind that the patterns observed within the CR module framework might not upscale straightforwardly to multitrophic networks due to non-trivial indirect interactions occurring in multispecies communities. In the main text, we study the evolution of the traits *m*_1_ and *f*_1_ separately, but we also tackle the co-evolution of the two traits in the supporting information (Fig. S2, Appendix S1.B.4).

## Results

### Warming induces diversity losses within trophic networks

Warming is responsible for considerable biodiversity losses within multitrophic networks. This is true with (Fig. 2d) or without evolution (Fig. 2a). Though diversity recovers fast when evolution is allowed, note that diversity collapses to half of its initial value (Fig. 2d). Long after the transient state, diversity stabilizes at smaller values than before warming for a vast majority of simulations, with and without evolution (Fig 2g&h). A warming of 8°C for instance leads to a significant loss of trait diversity (around 32% without evolution, 13% with evolution). For a mutation rate *μ* = 10^−2^, the statistical model fitted explains around two thirds of the observed variance, with warming explaining almost one third of it (Table S2.A). The model confirms the expected tendency revealed by Fig. 2g&h: stronger warming leads to higher diversity loss. The estimated coefficients governing the linear dependence of persistence on warming intensity are significantly negative without evolution (*Warning_Effect_* = −0.024, *p_value_* < 2.10^−16^) or with evolution (*Warning_Effect_ + Interaction_Effect_* = −0.007, *p_value_* < 10^−3^).

**Figure 2:**
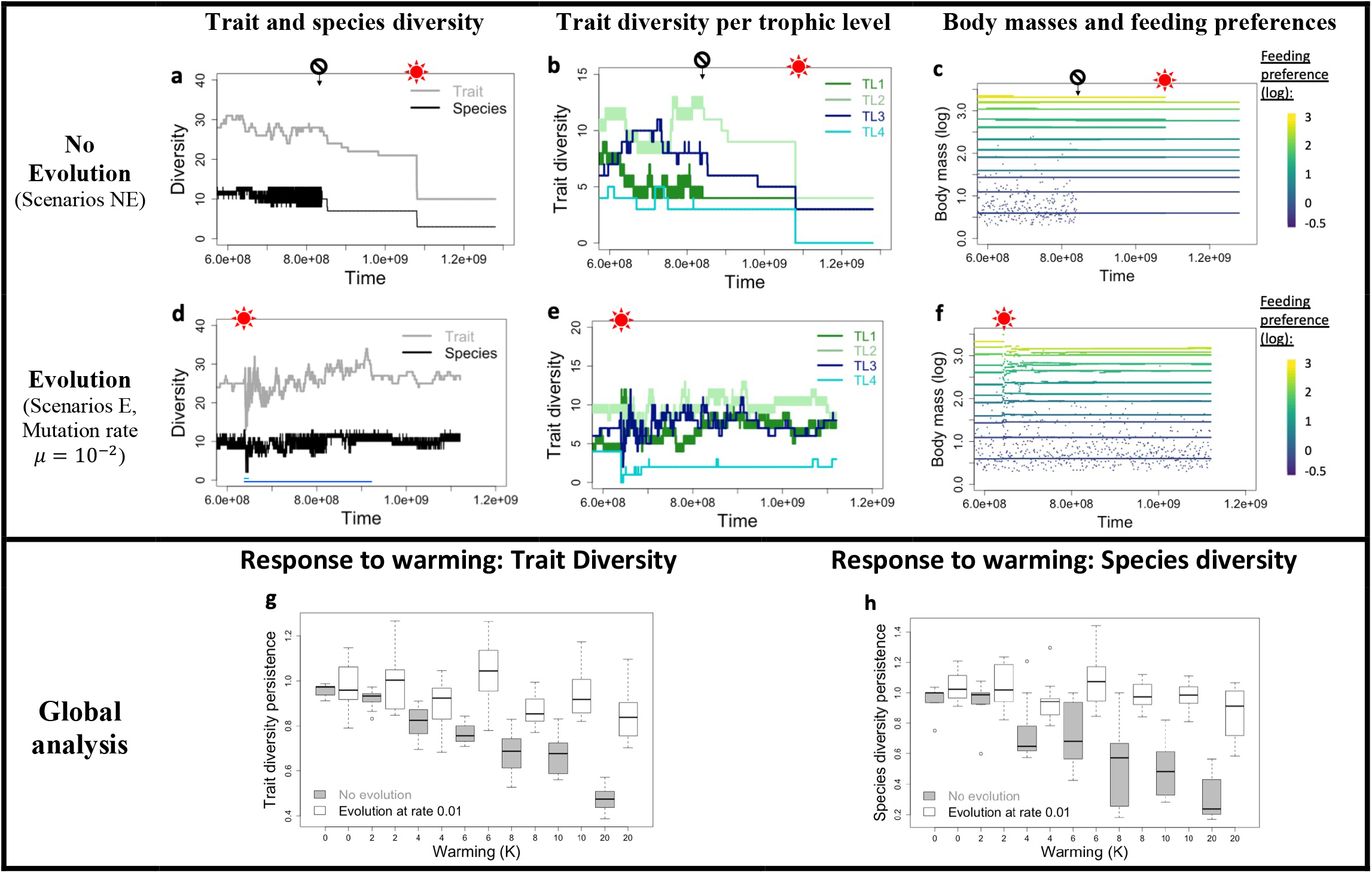
Diversity response to warming. **a-b-c:** Diversity and evolutionary dynamics of a simulation where evolution has been stopped (prohibition symbol indicates stopping time) before warming from 275 K to 295 K (sun symbol); **d-e-f:** Same outputs with ongoing evolution at warming. **g-h:** Boxplots of trait and species diversity persistence according to different warming intensities for scenarios with or without evolution. Each box corresponds to 10 independent replicas. See Table 1 for parameter values.

Diversity is lost because some consumer populations go extinct following the warming event. As temperature increases, less energy is available at the bottom of the trophic network (decline in basal resource’s carrying capacity *K*(*T*), equation 5), while the metabolic requirements increase (increase in respiration loss rate *d_i_*(*T*), equation 4). Despite an increase of plants productivity (basal resource’s growth rate *r*(*T*), equation 3), the ratio of ingestion to metabolic losses overall decreases with warming. In the CR framework, this decrease is driven by the ratio 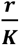:

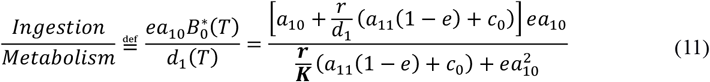

This ingestion ratio reaches the critical value of one at a critical temperature *T_c_*(*f*_1_), above which a consumer with feeding preference *f*_1_ cannot survive (Appendix S1.A). Ingestion is maximal when the consumer’s feeding preference matches exactly the basal resource phenotype. No consumer population can therefore survive above the limit temperature 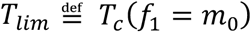 (equation 12, Appendix S1.A).

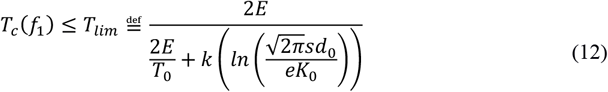

The limit temperature *T_lim_* being independent of the consumer’s traits, even fast evolution would not allow any consumer population to survive higher temperatures. Fig. 3 for instance shows how evolved body mass depends on temperature. Above *T_lim_* (≈ 316.4*K* here, grey area), no consumer phenotype is viable. Consumer survival depends on the intensity of warming: a warming from *T*_1_ to *T*_2_ would allow survival while a warming from *T*_1_ to *T*_3_ would lead to extinction. Note also that temperature does not impact evolved body mass in the CR module (analytical proof in appendix S1 B.2).

**Figure 3:**
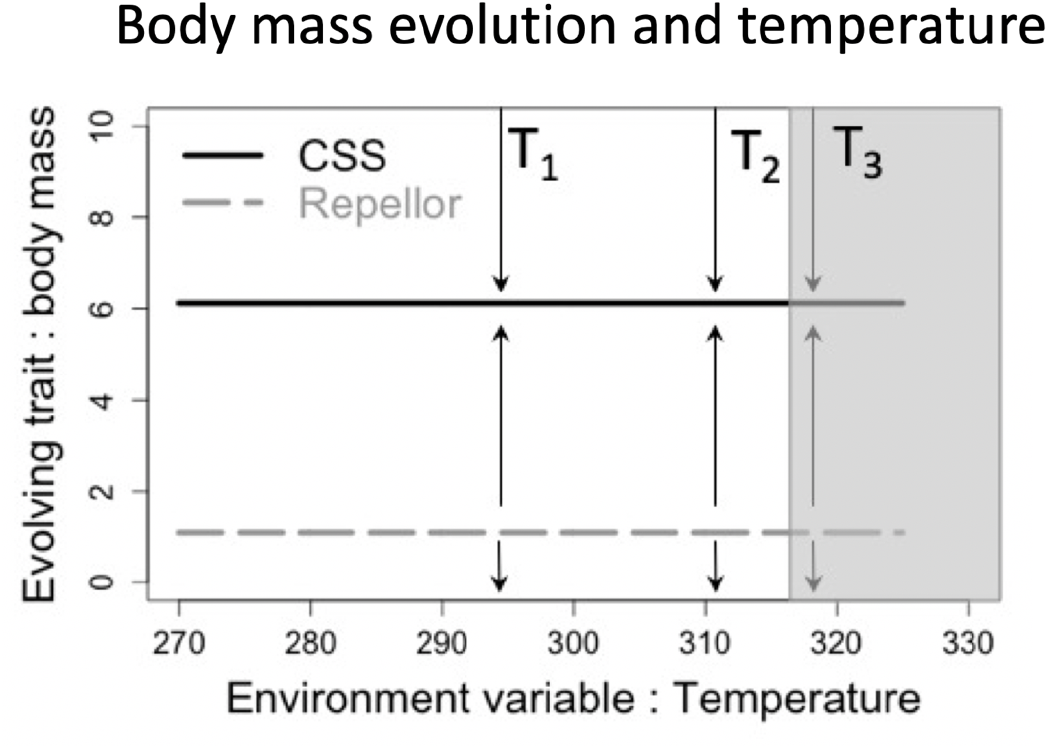
Body mass evolution and temperature (E3 diagram). Grey area cannot support the consumer. Arrows indicate the direction of evolutionary trajectories. Temperatures ***T*_1_, *T*_2_, *T*_3_** show different warming scenarios. Parameters: ***f*_1_ = m_0_**. Other parameters as in Table 1.

Warming can also reduce long-term diversity by changing the selection regime from disruptive to stabilizing. Within the CR framework, such a pattern is observed in the case of feeding preference evolution, for intermediate consumer to resource body mass ratios (Fig. 4, details in Appendix S1 B.3.c). As temperature increases, the convergent singularity changes from an invasible BP to a non-invasible CSS (Fig. 4a). Diversification at branching yields the emergence of a stable dimorphism of contrasting feeding preferences (≈ (*m*_0_, *m*_1_), Fig. 4c). The initial 2-level food chain (*t_a_*, Fig. 4d), consisting of a consumer population feeding equally on the basal resource and conspecifics (cannibalism), evolves into a 3-level food chain (*t_b_*, Fig. 4d). The intermediate level (Fig. 4d, 1) now primarily relies on basal resource consumption, making the upper trophic level (Fig. 4d, 2) viable despite being highly cannibalistic. As illustrated in Fig. 4b, a warming from *T*_1_ to *T*_2_ at branching (*t_a_*) completely changes the evolutionary dynamics. At *T*_2_, the loss of the upper trophic level leads to a 2-level food chain with a consumer population primarily feeding on the basal resource. Trait diversity is lost and will not recover as selection is stabilizing at *T*_2_ (CSS, Fig. 4a). Long-term warming-induced diversity losses can derive from a loss of diversification processes driven by warming.

**Fig. 4:**
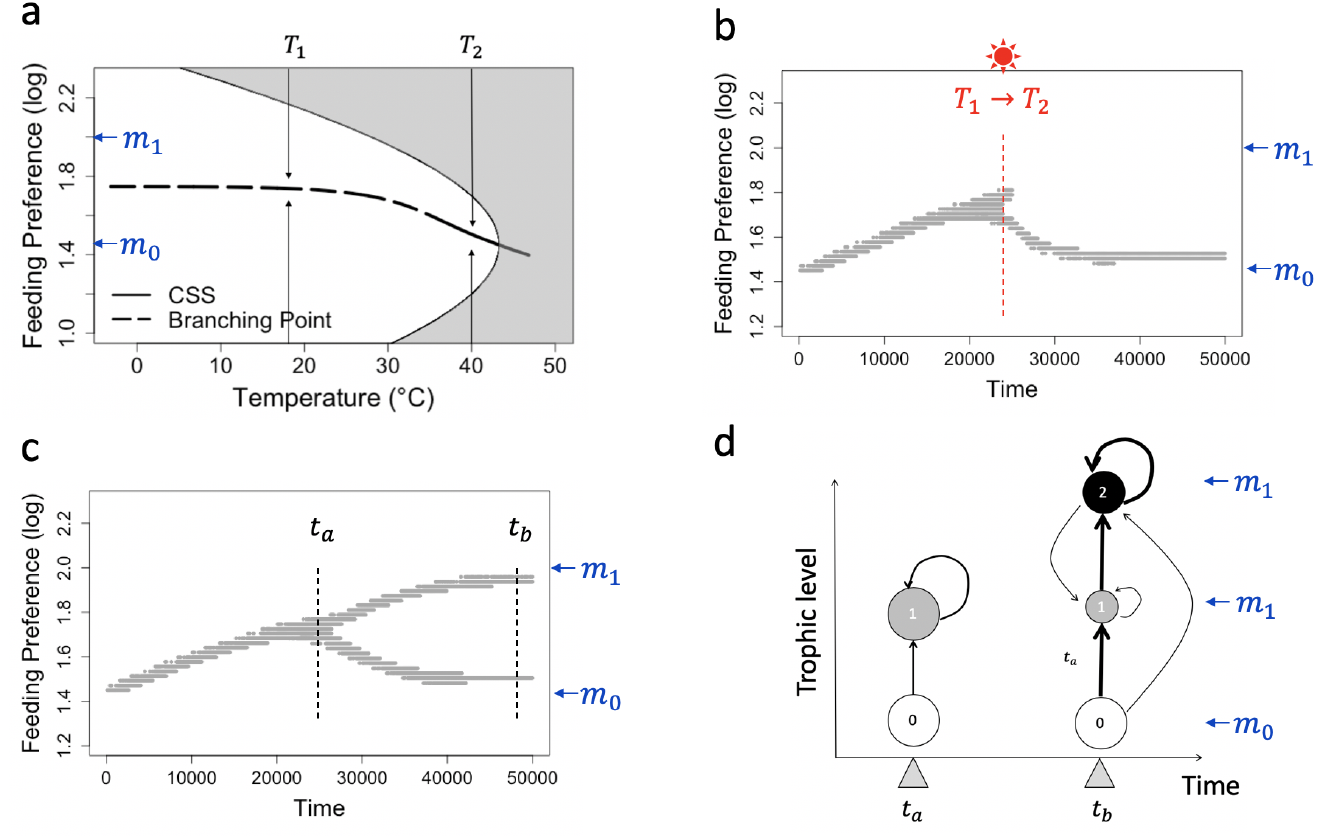
Warming switches the selection regime from disruptive to stabilizing. (CR module, evolution of feeding preference *f*_1_). *m*_1_ = 100, *m*_0_ = 28.2 **a.** Dependence of singularities on temperature. **b/c**. Evolutionary dynamics (mutation rate *μ* = 10^−2^, mutation steps ±5%) at temperature *T*_1_ (**c**, Branching Point) or for a warming from *T*_1_ to *T*_2_ occurring at branching (**b**, sun symbol). **d.** Schematic view of the trophic network before (*t_a_*) and long after branching (*t_b_*). Arrow thickness shows the intensity of trophic interactions. Circle size is proportional to the morph density. Other parameters are as in Table 1.

In our simulated networks, upper trophic levels, corresponding to consumers with high body masses and/or feeding on large prey (Fig. 2c&f), are most vulnerable to warming (Fig. 2b&e). This may be explained by three factors. Firstly, the analysis of the CR module reveals that a higher body mass is responsible for a sharper decrease of the ingestion ratio with warming (Fig. S1, Appendix S1 A.4). Secondly, in a multitrophic context, upper trophic levels suffer from accumulated warming-induced energy losses happening at lower trophic levels. Thirdly, a final reason for the vulnerability of upper trophic levels is their low population sizes. This reduces their evolvability, thus their ability to adapt to new environmental conditions. Warmed trophic networks are consequently flatter. Warming-induced losses of diversifying processes that yield the emergence of upper trophic levels, such as the branching point in Fig. 4, could also contribute to the pattern observed.

### Evolution can mitigate warming-induced diversity losses

Warming triggers diversity losses with or without evolution (Fig. 2a&d). However, recovery is possible when morphs can evolve (Fig. 2d), while it is not when evolution has been switched off (Fig. 2a). Such a recovery necessitates a considerable amount of time. In Fig. 2d, it took 200 000 mutation events/generations (light blue line) to reach pre-warming diversity levels and far more for diversity to be stable over time (2.8 10^6^ mutation events/generations, dark blue line). The diversity eventually maintained in the trophic network is significantly higher in scenarios with evolution (Ancovas, Table S2). The results are quantitatively illustrated on trait diversity for a mutation rate *μ* = 10^−2^ but remain consistent across the three mutation rates tested. Evolution has a positive effect on diversity persistence. This effect is both direct (*Evolution_Effect_* = 0.06, *p_value_* = 0.0204, *Var_explained_* = 28.4%) and indirect, increasing in importance with warming (*Interaction_Effect_* = 0.017, *p_value_* < 10^−8^, *Var_explained_* = 9.5%). This interactive effect is strong enough to totally offset the negative linear dependence of diversity persistence on warming in scenarios with evolution for the two other mutation rates tested (10^−1^, 10^−3^). In these cases, trait diversity recovers totally after a long transient state. Due to the interaction term, the stronger the warming, the higher the diversity maintained thanks to ongoing evolution across the trophic network (Fig 2g&h). It reaches a maximum of almost 40% at 20°C, with 48% of trait persistence in scenarios without evolution vs 85% in scenarios with evolution. Among evolutionary scenarios, we also tested the theoretically known association between faster evolution and better rescue by comparing diversity maintenance across mutation rates (ANCOVA). Counterintuitively, no significant difference was found (but see discussion).

The analysis of the CR module highlights two processes that potentially explain how evolution contributes to preserve the diversity in multitrophic networks: evolutionary rescue and an indirect mechanism we label “diversity-mediated buffering effect”. These processes, which are illustrated in the case of feeding preference *f*_1_ evolution, are explained in detail further below.

It can be shown that the consumer’s evolved feeding preference 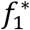 is necessarily between its own body mass (*m*_1_) and the body mass of the resource (*m*_0_) (Appendix S1 B.3). The study of feeding preference in this interval is thus sufficient. We define the consumer trait 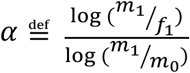 describing the proportion of resource consumption on the total consumer regime. Strictly equivalent to feeding preference evolution, it is more convenient to study the evolution of a that varies between two extreme scenarios: the consumer is essentially cannibalistic (*α* = 0) or primarily relies on the basal resource ingestion (*α* = 1). The analysis of the CR module reveals that four qualitative outcomes, corresponding to four ecological scenarios, are possible (Appendix S1 B.3.c, Table S1). These scenarios are illustrated in Fig. 5. They differ in two key features: (1) the consumer to resource body mass ratio 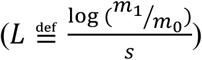 is either small (scenario A), intermediate (scenario B) or large; (2) for large ratios, whether the consumer body mass (*m*_1_) is small (scenario C) or big (scenario D). Note that dynamics presented earlier (Fig. 4a) correspond to scenario B (Fig. 5c).

**Figure 5:**
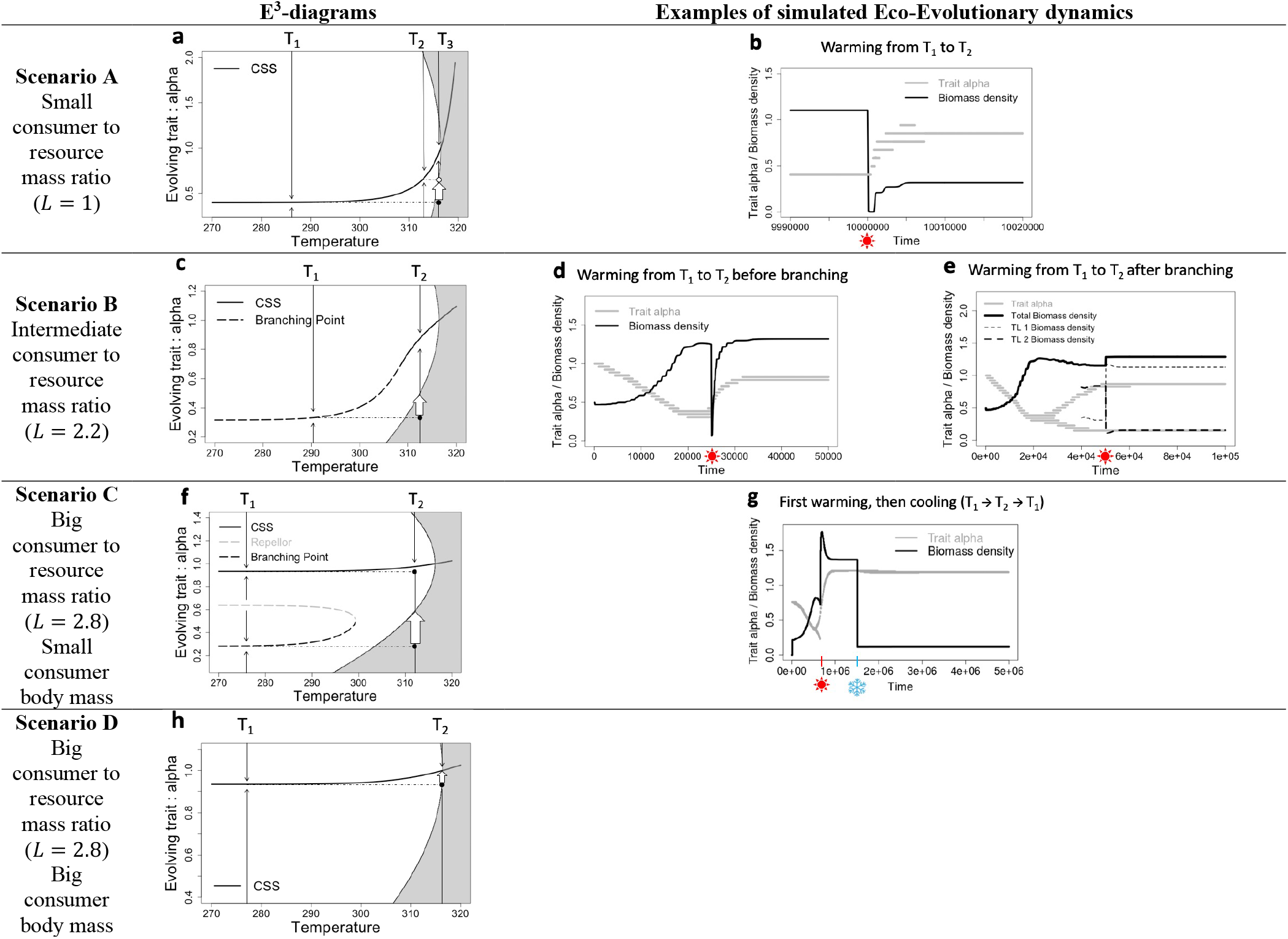
**a-c-f-h E^3^ diagrams** corresponding to scenarios A, B, C and D (Table S1). Grey areas are non-viability regions; the curves indicate the evolutionary singularities. Thin black arrows indicate the direction of evolution and the big white arrows indicate potential evolutionary rescues. **b-d-e Model simulations with evolutionary rescue occurring** (sun indicates warming time). Evolutionary rules: *μ* = 10^−2^ and small mutational steps: 5%*f*_1_. **g-** Model simulation with evolutionary hysteresis occurring (sun: warming; snowflake: cooling). Evolutionary rules: *μ* = 10^−2^ and small mutational steps: 1%*f*_1_. Parameter values are in Table 1.

Evolutionary rescue can never happen for temperatures above *T_lim_* (equation 12), but is always possible if the final temperature remains below *T_lim_* (Fig. 5). Consumers that adapt fast enough to the new conditions will avoid extinction, as figured by the white arrows in Fig. 5a-c-f-h. For instance, at small CR body mass ratios (scenario A, Fig. 5a), a warming from *T*_1_ to *T*_3_ leads the system to enter the non-viability area (black dot in grey area). The food chain can however persist if evolution is fast enough (white arrow), as larger values of *α* are selected, outside the non-viability area. Such values imply higher intakes from the basal resource relative to cannibalism that enable the maintenance of a sufficient ingestion rate despite the deteriorated environmental conditions. The selection of higher intake from the basal resource relative to cannibalism seems consistent across scenarios (Fig. 5). Simulations confirm the possibility of evolutionary rescue (Fig. 5b). The consumer population collapses, but if a evolves fast enough, the population recovers. Note that if warming is slow from *T*_1_ to *T*_2_ first and then from *T*_2_ to *T*_3_ (Fig. 5a), the trait would remain close to the CSS curve, and the population would never be threatened. Progressive warming decreases extinction risk, in agreement with theoretical (Gomulkiewicz & Holt, 1995) and empirical work (Bell & Gonzalez, 2009).

When evolution allows the diversification of the consumer niches, warming is potentially less threatening as the different occupied niches are unlikely to be simultaneously destroyed. We propose to label this mechanism “diversity-mediated buffering effect”. This can be observed in scenarios where branching leads to a stable dimorphism among consumers (scenarios B&C) so that a three-level food chain emerges (Fig. 4). We simulated the eco-evolutionary dynamics following a warming from *T*_1_ to *T*_2_ affecting either the two-level food chain (Fig 4d, just before branching at time *t_a_*) or the three-level food chain (Fig 4d, long after branching at time *t_b_*). While warming at *t_a_* largely threatens the population, the impact of warming is vastly reduced when temperature changes at *t_b_* (Fig. 5d vs 5e). Note that while the top trophic level biomass still suffers from the disturbance, the intermediate consumer phenotype actually benefits from warming (Fig 5e). Total consumer biomass increases. Interestingly, the selected intermediate consumer’s trait *α* pre-warming is close to the one selected under warmed conditions (Fig. 5e vs 5d).

Our simulated multitrophic networks emerge via successive branching events starting from a single consumer population initially. As a result, the consumer morphs occupy a wide diversity of niches (Fig. 2c&f) that ranges across 4 trophic levels (Fig. 2B&C). In other words, this “diversity-mediated buffering effect” is at play at the multitrophic network scale because multitrophic networks are the result of the consumer niche diversification.

### Evolution can also exacerbate the consequences of warming on diversity

While evolution often facilitates diversity maintenance in our simulated trophic networks, note that it cannot totally mitigate the negative effect of warming on diversity. Actually, in some cases we found that evolution can even exacerbate the negative consequences of warming-induced diversity losses. For instance, the analysis of the CR model suggests that warming can lead to eco-evolutionary tipping points that would severely depress persistence. Given scenario C, warming from *T*_1_ to *T*_2_ modifies the number of singularities (Fig. 5f). At *T*_1_, three singularities exist (CSS, Repellor and BP) while at *T*_2_ only the CSS remains. Consequently, starting from a resident consumer’s trait near branching, warming would be responsible for a reduction of diversity, as the system switches from selective pressures that allow stable polymorphism (BP) to a monomorphic situation (CSS). While similar to scenario B (Fig. 4), a major difference exists. Given the convergent properties of the CSS, decreasing temperature will not recover the initial diversity, as phenotypes remain at the selected CSS. This is a case of evolutionary hysteresis. Simulations confirm such dynamics (Fig. 5g). Starting near branching, we warm the system from *T*_1_ to *T*_2_ and observe the loss of the polymorphism. When the system is cooled back to *T*_1_, the system remains monomorphic. Here, the initial diversity lost to warming cannot be recovered by reducing temperature because of the eco-evolutionary constraints on consumer’s evolution. This result raises the possibility that diversity losses may be long-lasting and difficult to reverse, as a result of abrupt changes in selective regimes.

## Discussion

Our investigation into the effect of warming within multitrophic networks shows that warming is responsible for important diversity losses across food webs. While evolution helps to maintain biodiversity, it is certainly not sufficient to totally mitigate diversity losses. Evolution acts in at least two complementary ways. (1) By producing diversified ecological niches, as a result of disruptive selection, it leads to trophic networks that are more resistant to warming. (2) After warming, evolutionary rescue processes across the network lead to gradual and partial recovery. Evolution can however also exacerbate the negative consequences of warming, for instance by making them last longer (hysteresis), due to the crossing of eco-evolutionary tipping points. The consistency and coherence of the picture obtained by combining two complementary frameworks, with different scales and complexities, gives confidence on the robustness of our work. Our key results, as discussed in more detail below, have potential implications for the preservation of biodiversity in the context of current warming.

Warming induces important diversity losses within trophic networks. As experimentally observed (Rall et al., 2010), the ratio of ingestion to metabolic losses decreases because less energy is available at the bottom of the food web (primary producers) in a context of increasing metabolic expenditures. As a result, several consumer morphs go extinct. These extinctions are more likely to happen high in the food web. This result seems firmly grounded as several studies using a large diversity of approaches found a similar pattern (Petchey et al., 1999; Pounds, Fogden, & Campbell, 1999; Binzer et al., 2012). Potential explanations combine a sharper decrease of the ingestion ratio for morphs that are closer to their viable minimum; the bottom-up accumulation of deleterious consequences and the low evolvability associated with smaller population sizes and longer generation times. Warming also reduces diversity by altering evolutionary processes. Our CR framework shows that it can modify the selection regime from disruptive to stabilizing (scenario B&C). Altering diversity patterns through such modifications of selection regimes has deeper implications. Such effects are likely to last longer and be more difficult to reverse because of the additional constraints they entail (e.g. hysteresis). Our results support the idea that conservation ecology should focus more on preserving the processes facilitating and maintaining diversity rather than diversity patterns per se (Smith, Bruford, & Wayne, 1993).

Evolution partly mitigates warming-induced diversity losses within multitrophic networks, as shown by our statistical analysis (Table S2). Two complementary mechanisms are likely at play. First, evolution reduces diversity losses through a “diversity-mediated buffering effect”. This relies upon two observations: evolution (disruptive selection) leads to the diversification of the consumer niche (successive branching events) that allows the emergence of multitrophic networks; the greater the diversity of occupied niches at warming, the more robust is the food web because of the increased likelihood of some strategies being resistant (Fig. 5 d&e). This mechanism is akin to the “insurance hypothesis” proposed to explain the resilience of diverse systems (Yachi & Loreau, 1999). Second, evolution allows diversity to progressively recover through evolutionary rescue processes. In our CR framework, evolutionary rescue is indeed often possible when evolutionary changes in foraging strategies allow higher energy acquisition or when evolution of body sizes reduces energy requirements. Such changes in either body size or foraging strategy in response to warming have been documented in nature. The metanalysis of Daufresne et al. (2009) for instance shows a significant decrease in the size of ectothermic aquatic organisms in response to climate change. In the United Kingdom, Pateman et al. (2012) showed that the butterfly *Aricia agestis*, originally a specialist of *Helianthemum nummularium* as a larval host plant, has been able to widen its foraging strategy, allowing its expansion in the face of climate change. It now also largely uses *Geranium mole* which is more abundant in warmer climates.

In our simulated multitrophic networks, evolutionary rescue likely happens for low and intermediate trophic levels (Fig 2f), which evolve faster due to larger population sizes. It is facilitated by occasional mutations with large phenotypic effects. Top trophic levels recover eventually from intermediate trophic level populations that evolve higher body masses and feeding preferences until occupying the niche freed by the extinction of top trophic levels. The recovery of the network’s diversity is however partial as lower biomass is available from primary production. Surprisingly, we found that higher mutation rates, associated with faster evolution, are not systematically associated with higher diversity persistence. A possible explanation is that, in addition to the intensity of selection, network recovery depends on the variability on which selection can act. Eventually, more frequent mutations do not yield more variability, but redundant phenotypes. This idea is in line with experimental results by Fugère et al. (2020) which clearly highlight that past a certain genetic variability, no improvement is observed in the rescue process. The work of Fugère et al. (2020) additionally illustrates how experimental evolution in microcosms or mesocosms offer promising opportunities for the critical empirical investigations into evolutionary rescue, especially at the multitrophic scale.

Evolution can however exacerbate the negative consequences of warming. Evolutionary hysteresis for instance makes them difficult to reverse, lasting longer. It implies strong and possibly irreversible shifts between alternative states corresponding to tipping points (Suding & Hobbs, 2009). In line with previous studies (Dakos et al., 2019), our work emphasizes the importance of considering both evolution and ecological dynamics to understand tipping points, especially in the context of global changes where selective pressures are likely strong. Observed in a simple food chain (CR, Fig. 5g), eco-evolutionary tipping points might also exist within complex multitrophic networks but such an investigation goes beyond the scope of the present paper. We simply note here that the transient state’s considerable duration following warming (e.g. Fig. 2d) potentially offers opportunities for hysteresis arising from the coevolution of traits and interactions we did not consider here. What is more, the overall vulnerability of the small populations during the transient state raises additional challenges. Demographic stochasticity or drift could dampen or impede progressive recovery. Important ecosystem services are likely to be degraded for a great period on a human timescale. However, we note that a recent study of the Cretaceous-Paleogene mass extinction suggests that functional recovery may happen much faster than diversity recovery (Alvarez et al., 2019).

Our parametrization relies, as much as possible, on available empirical data. Yet, our models are of course a crude simplification focused on a question, making many simplifying assumptions. For instance, we assume the carrying capacity of the basal resource to decrease exponentially with temperature based on the empirical data analyzed by Fussmann et al. (2014). This relationship is however debated and likely context-dependent, varying with the explicit limiting nutrient dynamics (Uszko et al., 2017). Moreover, while there is evidence for an increase of attack rates with temperature (Rall et al., 2010), available data suggest this effect is rather weak (Binzer et al., 2012; Rall et al., 2012). A hump-shaped relationship also seems more realistic when considering wide temperature ranges (Englund et al., 2011). Because it is not the focus of the present work (but see Weinbach et al., 2017 for an analysis of this question on the present model), the lack of a large scientific consensus led us to choose constant attack rates. Likewise, conversion efficiencies show large variations in nature and may depend on trophic positions (Lindeman, 1942). Herbivores (TL1) feeding on primary producers (TL0) typically have lower conversion efficiencies (Yodzis & Innes, 1992). We investigated the robustness of our results by varying this parameter in the CR module (Appendix S1.B.3). At lower efficiencies, we no longer observe evolutionary hysteresis, but typically have evolutionary rescue happening. Hysteresis is thus associated with high conversion efficiencies characteristic of higher trophic level (carnivores, Yodzis & Innes, 1992), which can be interpreted as additional evidence that higher trophic levels are more vulnerable to warming (Binzer et al., 2012). Overall, while lower conversion efficiencies certainly imply lower energy flows resulting in more vulnerable networks, there is no obvious reason to think the positive effects of evolution on diversity maintenance would change.

The concept of evolutionary rescue, originally formulated within a monospecific framework (Gomulkiewicz & Holt, 1995b), seems to extend to the community scale as our work suggests that multitrophic networks confronted with warming perform better when evolution is at play (Ferriere & Legendre, 2012). Among these processes, the evolutionary rescue of low and intermediate trophic levels, that facilitates the recovery of higher trophic levels, is key. The diversification of ecological niches ensuing from disruptive selection is equally important as our work unravels a diversity-mediated buffering effect. As a result, all measures favoring evolvability, such as spatial or temporal variability or the presence of microhabitat, are likely to make the community more resistant to warming. Likewise, all measures targeting the key factors for evolutionary rescue, as presented in the review by Carlson et al. (2014), should favor trophic networks’ persistence and resilience.

Yet, evolutionary processes are not sufficient to preserve biodiversity in the context of global change. Indeed, our work highlights important diversity losses for considerable amounts of time before a partial recovery can potentially happen. Specifically, the transient collapse may severely affect ecosystem services sustained by ecological networks, with large impacts from a management point of view. What is more, several mechanisms we do not consider here could dampen or even prevent the recovery. Low population size may induce adverse ecological (demographic stochasticity, Allee effects) or evolutionary effects (genetic drift) impeding persistence. We also did not model the evolution of primary producers under warming, which may mostly affect other traits than body size (Parmesan, 2006). Empirical evidence highlights for instance temporal mismatches between plants and herbivores resulting from heterogenous phenological shifts in response to climate warming (Visser & Gienapp, 2019). The consecutive energy losses further threaten the maintenance of diversity within multitrophic networks.

In summary, while evolution has a positive effect on biodiversity maintenance within trophic networks confronted with warming, the impact of warming is nevertheless expected to be dramatic and long lasting, with severe consequences for human populations. Conservation and biodiversity management policies should better integrate evolutionary components to properly address the issues raised by global change.

## Supporting Information

Additional Supporting Information may be found in the online version of this article:

**Table S1.** Warming impacts the CR evolutionary dynamics: the 4 possible scenarios

**Table S2.** Diversity persistence in the complex multitrophic networks: statistical analysis summary

**Fig. S1.** The decrease of the ingestion ratio with temperature depends on body mass

**Fig. S2.** Consumer Resource module: the two-trait co-evolution scenario

**Appendix S1.** Eco-evolutionary dynamics of the Consumer-Resource module

**Appendix S2.** Complex multitrophic network model

**Appendix S3.** Simulation code

## Acknowledgement

This work was funded by the National Research Agency of France (ANR) as part of the ARSENIC project (ANR grant no. 14-CE02-0012). We also would like to thank the HPCave computational facilities of Sorbonne Université on which the simulations were run.

## Author’s contributions statement

NL, KTA and YY conceived the ideas and designed the methodology. The analysis was conducted by AW (body mass evolution) and YY (feeding preference evolution and coevolution). As for the simulated complex multitrophic networks, KTA developed the code, which was later adapted by YY to do the simulations. Results analysis was done by NL, KTA and YY. YY wrote the first draft of the manuscript that has been subsequently reviewed and edited by NL and KTA. All authors contributed critically to the drafts and gave final approval for publication.

## Data Accessibility Statement

The code used for the simulations (programming language C) is available in the supporting information (Appendix S3).

## Supporting Information

## Appendix S1: Eco-evolutionary dynamics of the Consumer-Resource module

This document is dedicated to present the analytical work supporting the results presented in the main document on the CR module. The main equations (main document) are indexed as in the main document while new important ones are indexed by a letter, in order to avoid confusion. The same applies for figures or tables. This document also serves to present some complementary side results we feel improve the understanding of our work.

### A. Ecological dynamics

Population dynamics are given by the following equations:

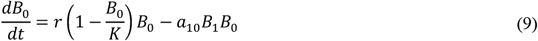

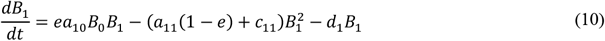

#### 1. Ecological equilibria

Ecological equilibria are given by resolving 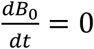 and 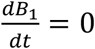 which leads to 4 possible solutions:

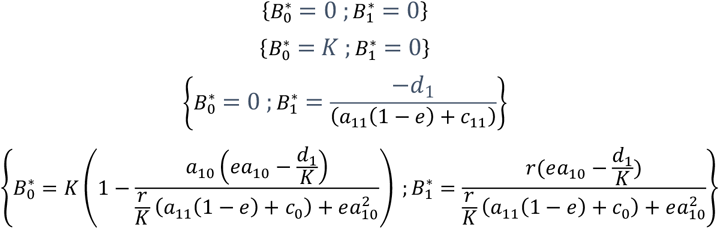

The third equilibrium is not reachable because 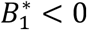. The fourth equilibrium is called the coexistence equilibrium.

#### 2. Ingestion ratio and coexistence

The coexistence equilibrium is possible when 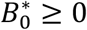 and 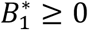.

Fist, we demonstrate: 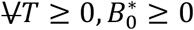 at the coexistence equilibrium.

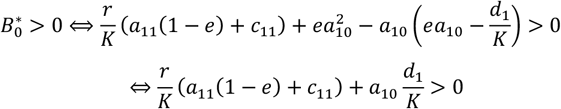

The last relation is always true which ends the proof.

The flux of biomass providing energy from resource to consumer 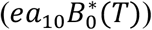 divided by the flux of biomass lost due to metabolic expenditures (*d*_1_(*T*)) corresponds to the ingestion ratio *IR*.

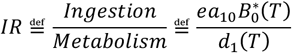

But

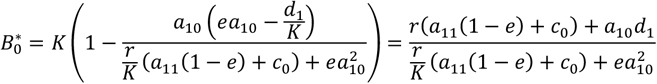

So that:

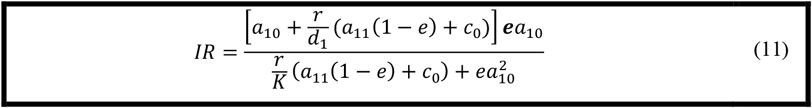

Now, we show that the consumer population is viable if and only if its ingestion rate is above one.

We have:

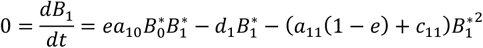

Which implies:

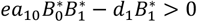

Now:

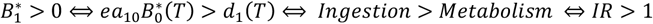

#### 3. Critical and limit temperature

We now prove that the ingestion ratio reaches a value of one at a critical temperature above which, according to the previous section, no consumer population can survive. For convenience, we introduce the following notations:

We define *M*_0_, *M*_1_, *F*_1_ by 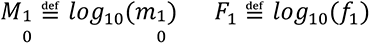

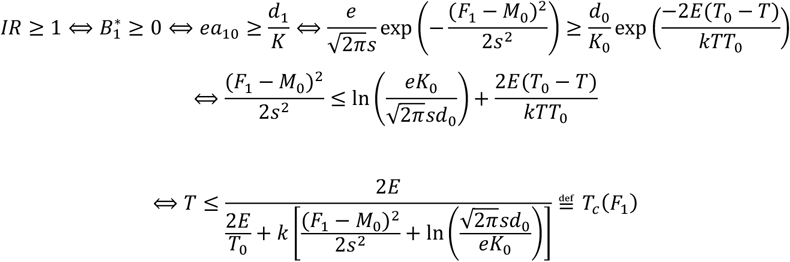

*T_c_*(*F*_1_) is maximal when its denominator is minimal, that is to say when *F*_1_ = *M*_0_. Note that this is the parametrization used in the case of body mass evolution, feeding center being fixed. It leads to equation (12):

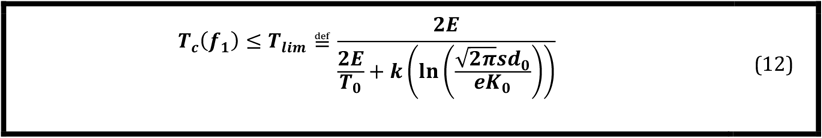

With the standard parameter values presented in table 1, *T_lim_* ≈ 316.4 K.

In the case of feeding preference evolution, the convenient way to express the condition for coexistence 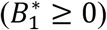, is the following (used for the viability areas in the E3 diagrams **Fig. 3**).

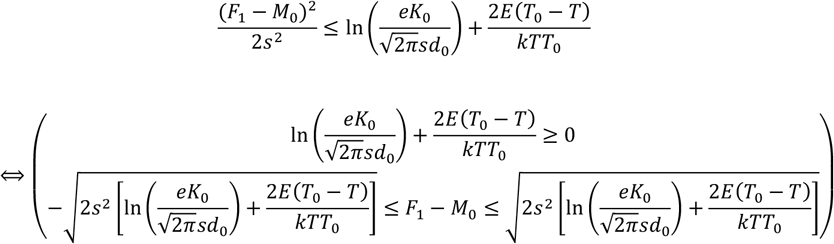

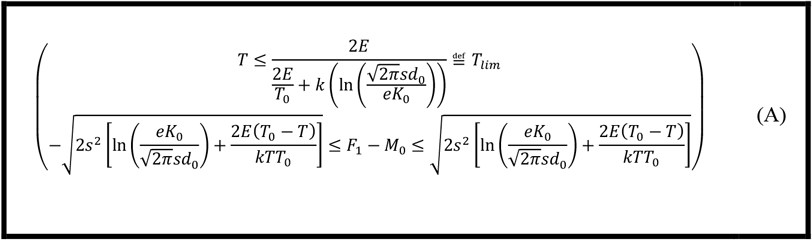

#### 4. Ingestion ratio and body mass

We have:

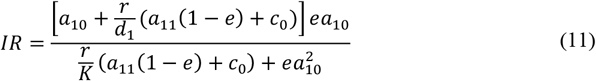

The only remaining dependence of temperature is due to the relative productivity 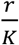 of the pool of plants forming the basal resource. It increases exponentially with temperature 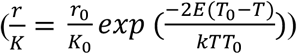 and explains that the ingestion ratio decreases with temperature. It is interesting to investigate this relationship for different consumer body masses *m*_1_ as bigger body masses are expected as one goes up the trophic network. **Fig. S1** shows that the ingestion ratio exhibits a sharper decrease with temperature as the consumer body mass increases.

**Figure S1:**
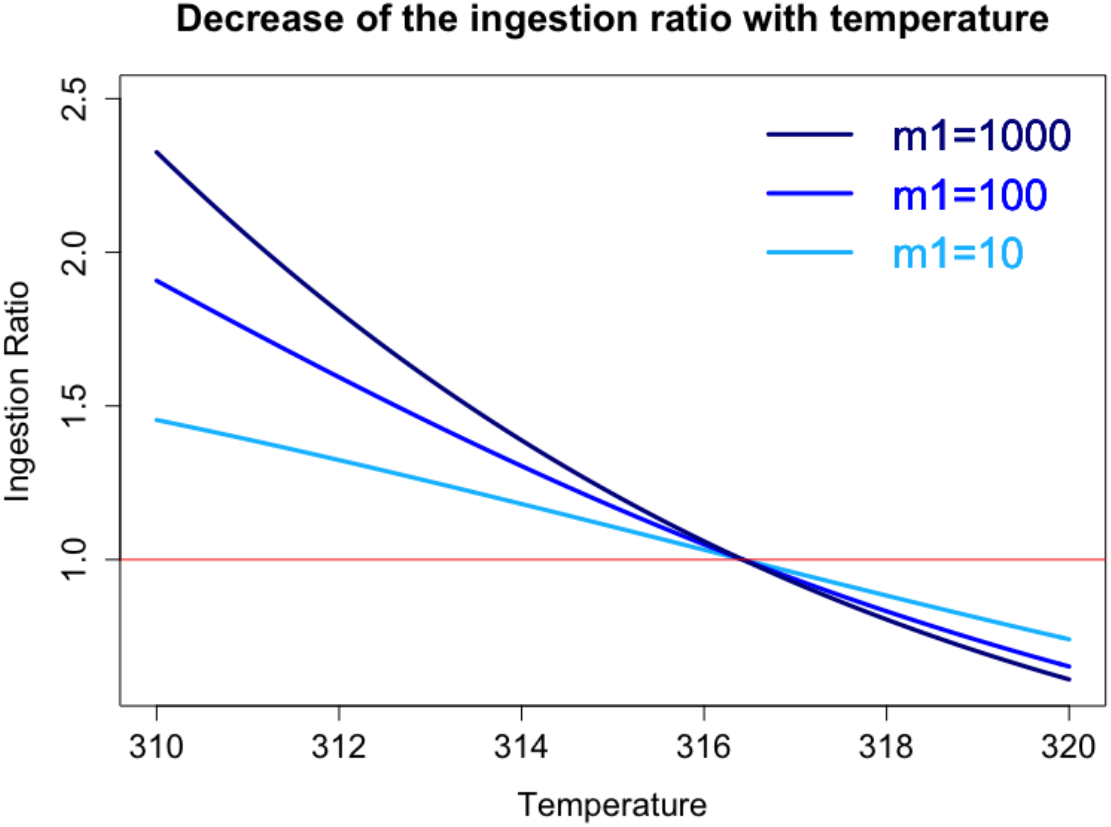
The ingestion ratio (equation (11)) decreases with temperature. Values above one (resp. below) indicate a net energy gain (resp. loss). The bigger the consumer body mass *m*_1_, the stronger the decrease. Note that the critical temperature *T_c_*(*f*_1_) above which a consumer with feeding preference *f*_1_ cannot survive is independent of its body mass *m*_1_. Here, we chose *f*_1_ = *m*_0_ = 1. All other parameters as in **Table 1.**

This result indicates that upper trophic levels, that are associated with bigger sizes (**Fig. 2c&f**), are likely to suffer more from warming than lower trophic levels. We would like to point out that the ranking of ingestion ratios with body masses observed in the CR module (*IR*(*m*_1_ = 1000) > *IR*(*m*_1_ = 100) > *IR*(*m*_1_ = 10)) do not contradict the fact that upper trophic levels are likely to be initially closer to their critical ingestion ratio. Indeed, the argument relies on the trophic distance from the basal resource, each trophic interaction being associated with losses (*e* < 1). Here (**Fig. S1**), the three consumer populations differ in body mass but have the same trophic level (= 1).

### B. Evolutionary dynamics

#### 1. The adaptive dynamics framework

In the case of body mass (*m*_1_) evolution, the invasion fitness 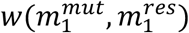 of a rare mutant corresponds to its relative growth rate in the resident population:

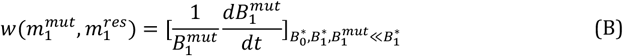

The sequence of trait substitutions describes the evolutionary dynamics of the system and can be approximated by the canonical equation (Dieckmann & Law, 1996) that links the trait dynamics to the selection gradient:

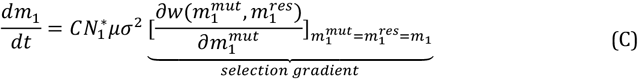

where 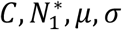 are respectively a homogenizing constant, the equilibrium resident population density, the mutation rate and the amplitude of mutations. The zeros of equation (C) are evolutionary singularities, 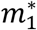, satisfying:

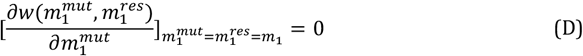

The convergence and invasibility properties of an evolutionary singularity are determined via the second derivatives of the invasion fitness function as detailed in section B.2.d.

#### 2. How does the selected body mass depend on temperature?

Reminder: we define 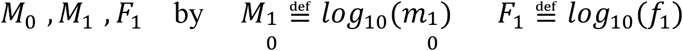

##### a. Fitness of invasion

Here, we expose the whole approach to determine the invasion fitness function in the adaptive dynamics’ framework:

We consider a rare mutant 1’ appearing in the resident population 1, given the population dynamics equations, we have:

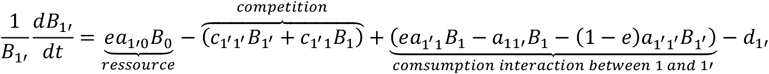

Given 1 and 1’ have the same feeding preference, we have *c*_1′,1′_, = *c*_1′1_ = *c*_0_ and *a*_11′_ = *a*_1′1′_. Also, *B*_1′_ ≪ *B*_1_ (rare mutant). Hence: *c*_1′1′_*B*_1′_ ≪ *c*_1′1′_*B*_1_ and (1 − *e*)*a*_1′1′_*B*_1′_ ≪ *a*_11′_*B*_1_ All in all, we deduce:

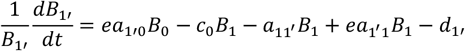

Which leads to:

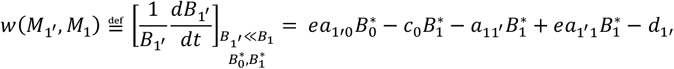

##### b. Evolutionary singularities

We first need to determine the derivative of the fitness invasion function with respect to the mutant body mass (first variable):

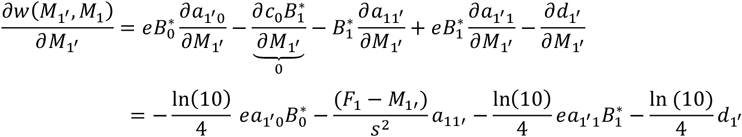

Thus:

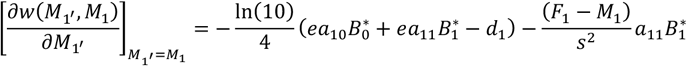

At ecological equilibrium 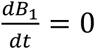 gives 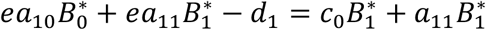 and hence:

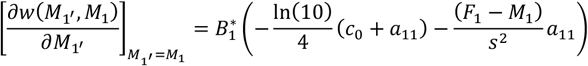

Evolutionary singularities occur when the selection gradient vanishes:

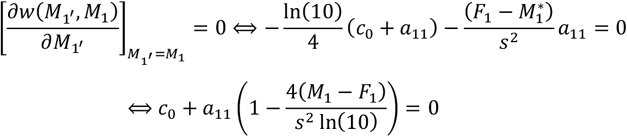

Noting explicitly the dependence of *a*_11_ on *M*_1_, we deduce evolutionary singularities 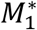 verify:

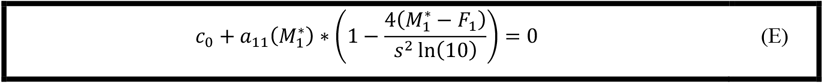

Temperature does not influence any parameter of equation (E) indicating that warming has no effect on the evolutionary dynamics of body mass here.

#### 3. How does the selected feeding preference depend on temperature?

##### a. Fitness of invasion

Here, the evolving trait is *f*_1_ or equivalently *F*_1_. As we did before, we consider the appearance of a new morph 1’ issued from 1 by a random small mutation.

Given our model, its biomass follows the equation (note that *d*_1_ = *d*_1′_):

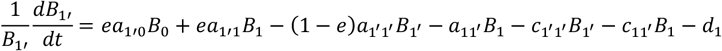

Using *B*_1′_ ≪ *B*_1_ and *F*_1_ ≈ *F*_1′_(small mutation hypothesis), we find:

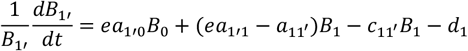

When considered at ecological equilibrium, the previous expression gives the invasion fitness function:

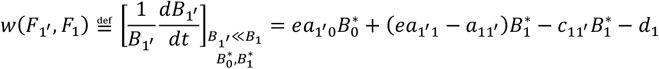

##### b. Evolutionary singularities

When the selection gradient vanishes:

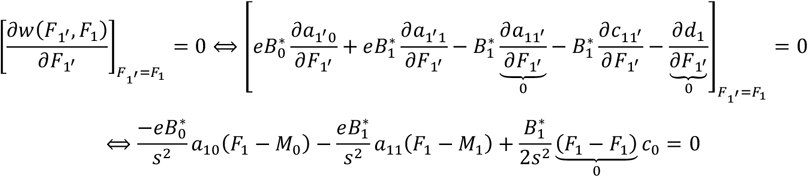

So that evolutionary singularities 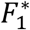 satisfy:

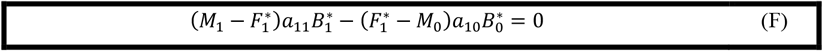

###### Proof of the existence of at least one singularity

Here, we prove that equation (F) has at least one solution and that all solutions 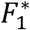 verify: 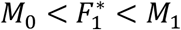. For convenience we define:

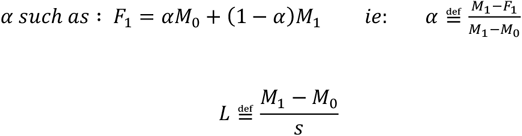

Equation (F) can be rewritten with these new notations, by noting that *M*_1_ − *M*_0_ (hypothesis of the model):

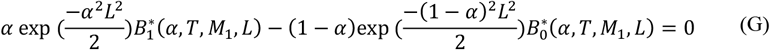

We define:

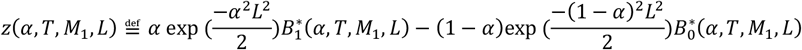

We have, *z*(0, *T, M*_1_, *L*) < 0, *z*(1, *T, M*_1_, *L*) > 0 and, given *T, M*_1_, *L z_α_*: *α → z*(*α, T, M*_1_, *L*) is a continuous function. So, by the Intermediate Value Theorem, there exists *α*\* such that *z_α_*(*α*\*) = *z*(*α, T, M*_1_, *L*) = 0. This is equivalent to say that equation (D) or (16) has at least one solution. Moreover, under coexistence (i.e. 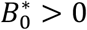 and 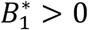), if *α* ≤ 0 *then z*(*α, T, M*_1_, *L*) < 0 and if *α*\* < 1 *then z*(*α, T, M*_1_, *L*) > 0 so that we have necessarily 0 < *α*\* < 1. This is equivalent to 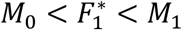. This ends the proof.

###### c. Effect of warming on feeding preference evolution and emergence of 4 scenarios

Since 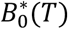 and 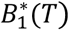 do depend on the temperature, equation (F) indicates temperature has an impact on the singularities and consequently on evolutionary dynamics. In order to go further, we rewrite (F) replacing 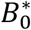 and 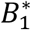 by their explicit expressions and isolating the temperature from the evolutionary singularity:

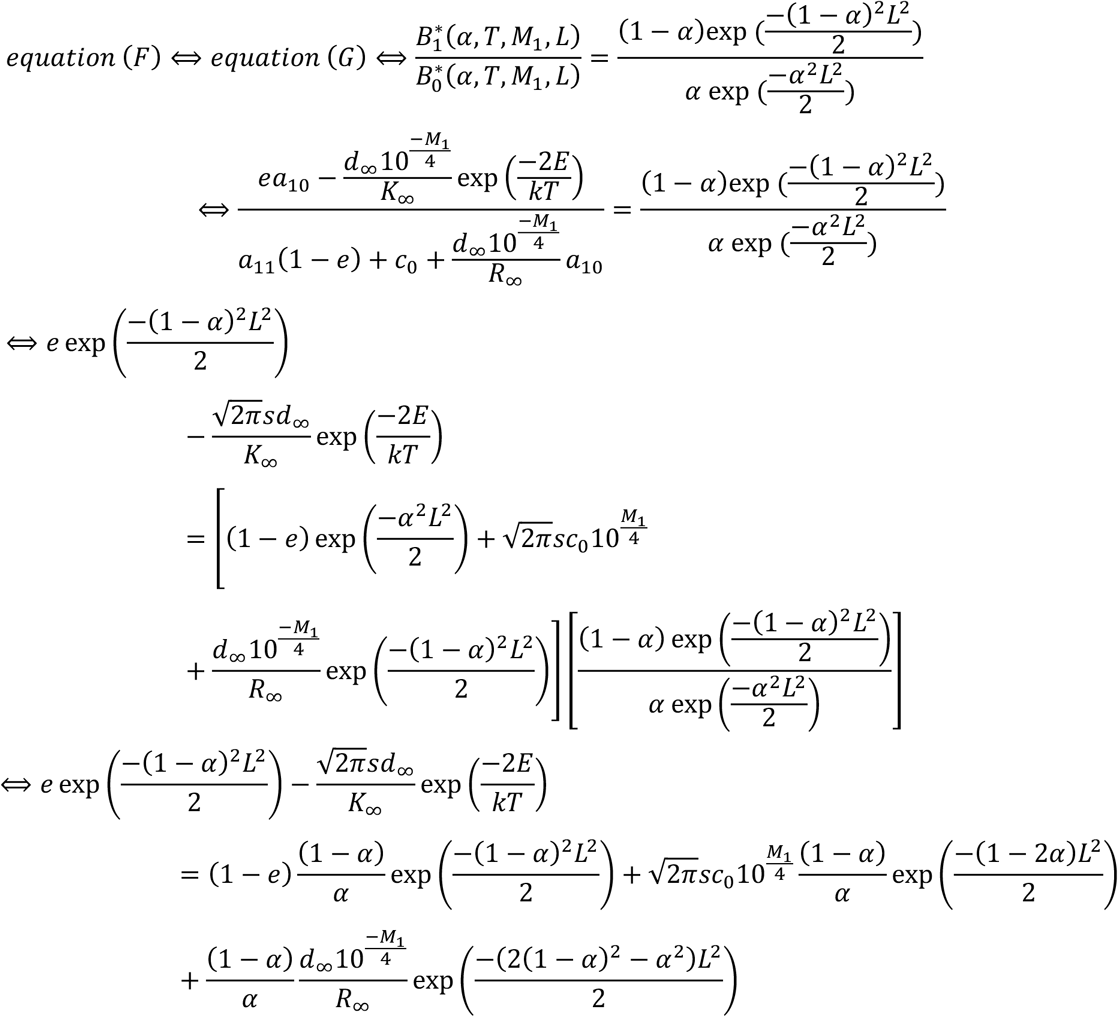

We define 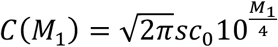 and have the equivalence with:

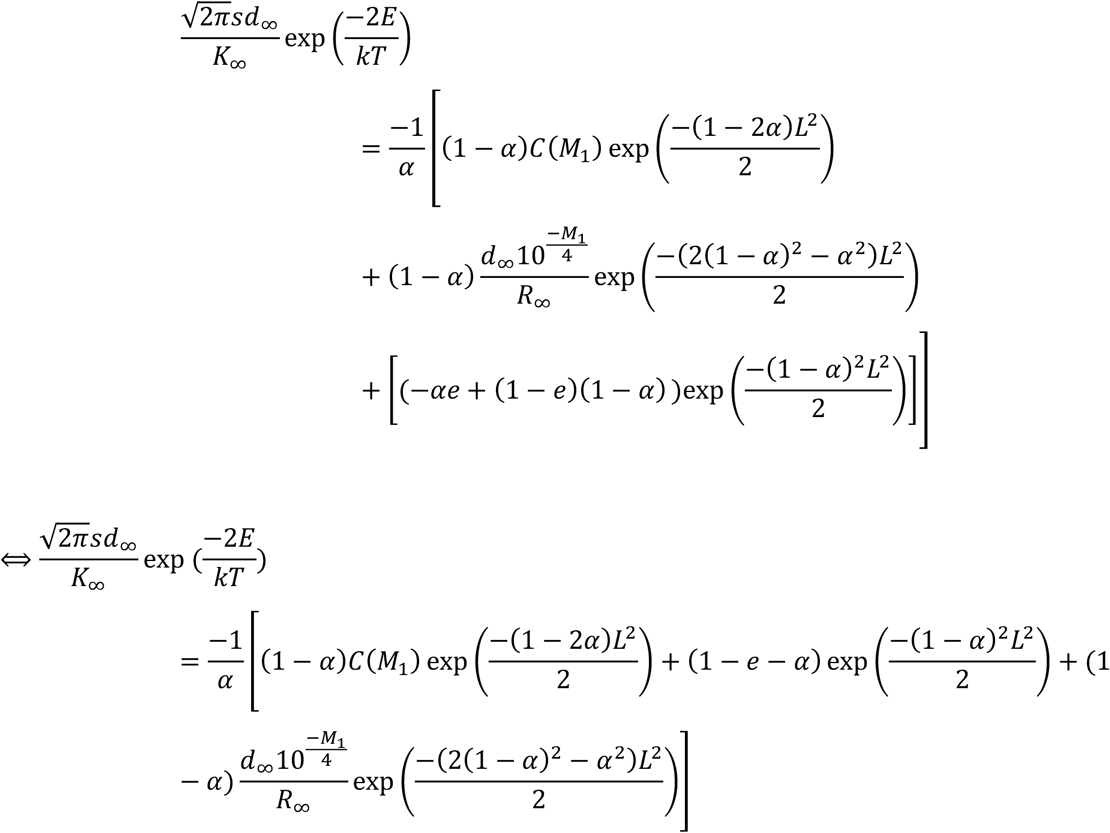

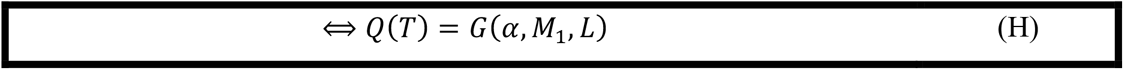

Where the functions *Q*(*T*) and *G*(*α, M*_1_, *L*) are defined by:

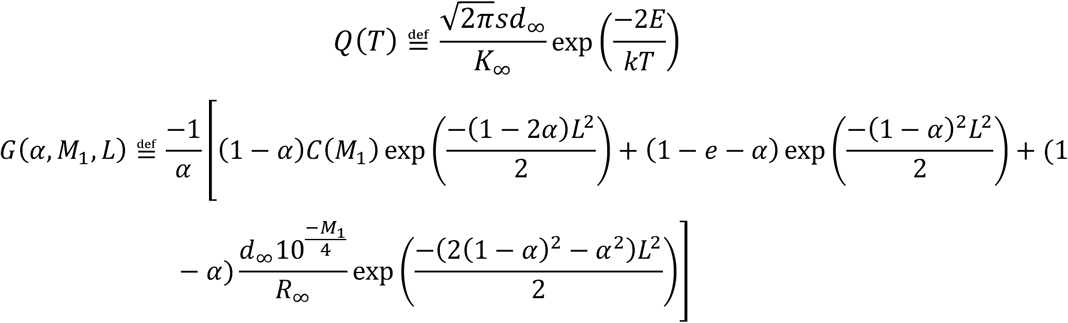

To investigate the effect of temperature on the evolutionary dynamics, we analyze the functions:

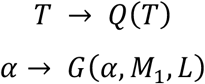

Q captures the temperature component of selection while G captures the biotic components of selection that depend on the consumer feeding preference (α), body mass (*M*_1_) and consumer to resource body mass ratio (L).

###### Study of Q as a function of T

*Q* is increasing with the temperature *T* and has the following variation table:

**Table A:**
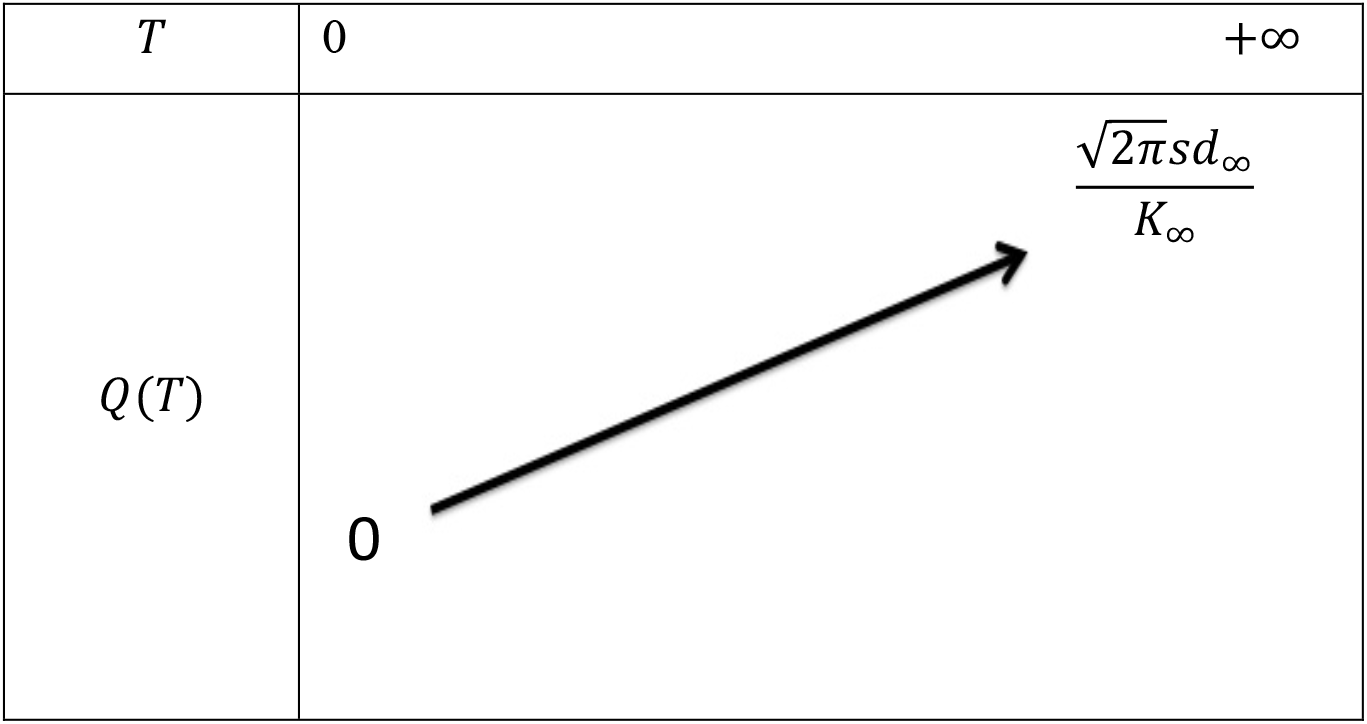
Table of variation of ***Q***(***T***). Note that ***Q***(***T***) is always positive.

###### Study of G as a function of α, M_1_ and L seen as parameters

Analytically we cannot determine all the possible behaviors of G. Thus, we used the software Python to investigate its variation. **Fig. B** shows the shape of G for different values of Consumer to Resource (CR) body mass ratios.

For small CR body mass ratios, *G* is increasing with a (**Fig. B a&d**). For big CR body mass ratios, G is not monotonous anymore (**Fig. B c&f**). There is a value of the parameter *L, M_1_*-dependant, we refer to as *L^c^*(*M*_1_), around which the switch of behavior from monotonous to not monotonous occurs. We can show by studying the derivative 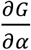 that 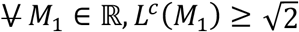. Its value is around 2.2 for the values of *M*_1_ we investigated (*M*_1_ = 0 and *M*_1_ = 2). In addition, just before the change of behavior occurs, the curvature of *G* changes (**Fig. B b&e**) meaning its second derivative switches sign. This can have consequences on the nature of the singularities as this nature depends on the second derivatives of the fitness gradient. This led us to consider scenarios with an intermediate CR body masses ratio.

**Fig. B:**
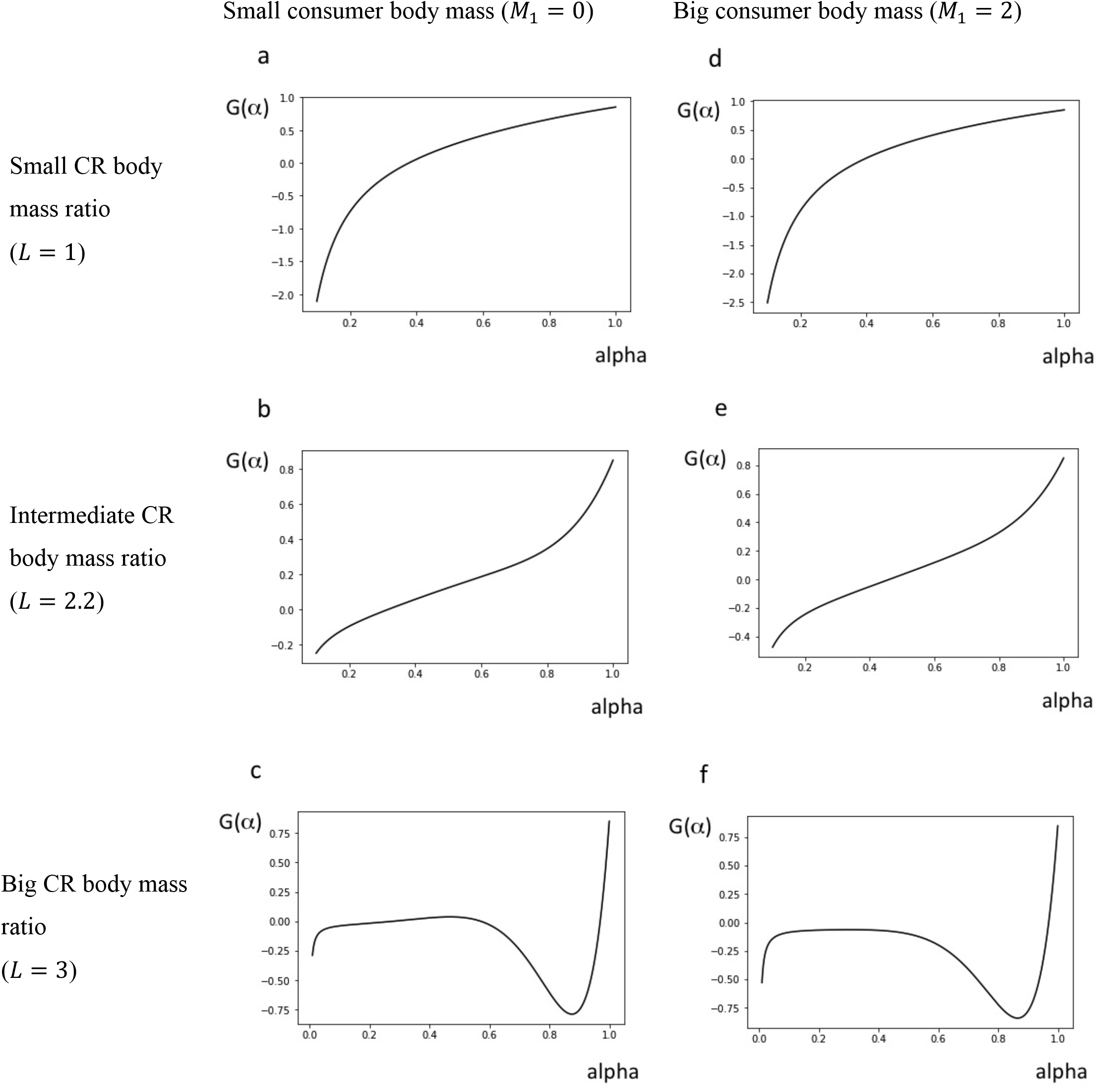
Plots of ***G***(***α***) for various values of ***L*** and ***M*_1_**.

From the previous analysis, we deduce the following table of variation for *G*:

**Table B:**
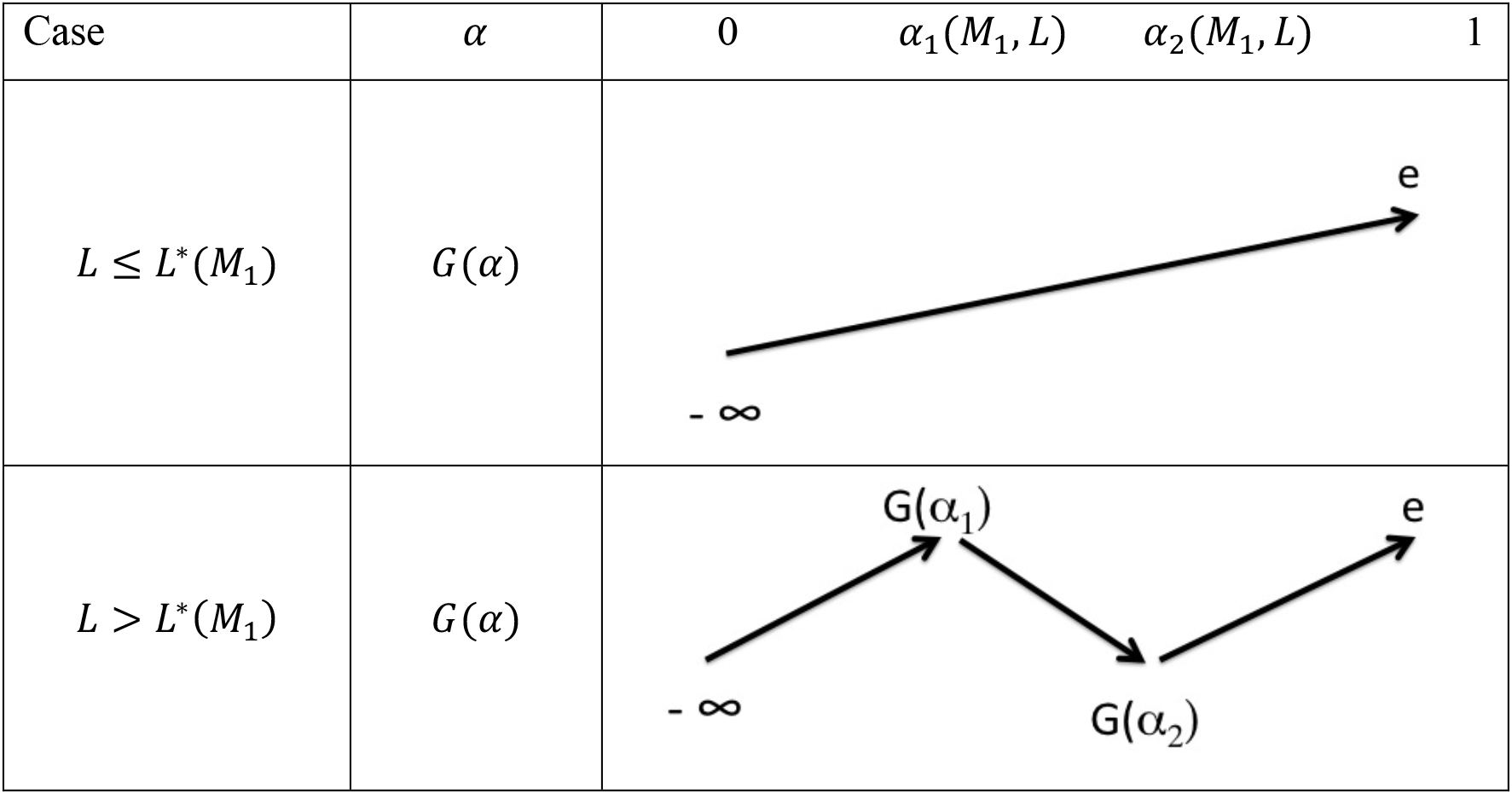
Variation table of function ***G*** according to the value of ***L***

If the consumer to resource body mass ratio is small, G is strictly increasing with a so that there is necessarily only one singularity.

If the ratio between consumer and resource body mass is big, the behavior of *G* indicates the existence of up to three singularities. The coexistence condition on temperature (*T_lim_*) implies *Q*(*T*) < *e* so that it is impossible to have two singularities (except one degenerate case) Hence, in this case, there is one or three singularities, depending on *G*(*α*_1_ (*M*_1_, *L*)), *G* (*α*_2_ (*M*_1_, *L*)) and *T*.

Finally, while there is no qualitative difference between the case of a small consumer body mass and a big one when the CR body mass ratio is either small (**Fig. A a&d**) or intermediate (**Fig. A b&e**), there is one in the case it is big. In that case, when the consumer body mass is small, *G*(*α*_1_) > 0 (**Fig. A c**), while when the consumer body mass is big, *G*(*α*_1_) < 0 (**Fig. A f**). This is important since *Q*(*T*) is always positive (see Table A). This means there can potentially be 3 singularities in the case of a small consumer body mass while there is only one in the case of a big consumer body mass.

All this analysis of equation (H) leads to distinguish 4 scenarios that differ in the impact of warming on feeding preference evolution: small CR body mass ratio **(scenario A)**, intermediate CR body mass ratio **(scenario B)**, big CR body mass ratio and small consumer body mass **(scenario C)**, big CR body mass ratio and big consumer body mass **(scenario D)**. These scenarios emerge through the technical analysis of the classical equation of adaptive dynamics (equation D) but are ecologically consistent as they differ in parameters known to be important for trophic interactions, namely body masses and body mass ratios 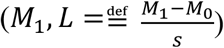. These scenarios are presented in the following **table S1**, with their ecological meaning and evolutionary dynamics.

###### Warming impacts the CR evolutionary dynamics: the 4 possible scenarios

**Table S1:**
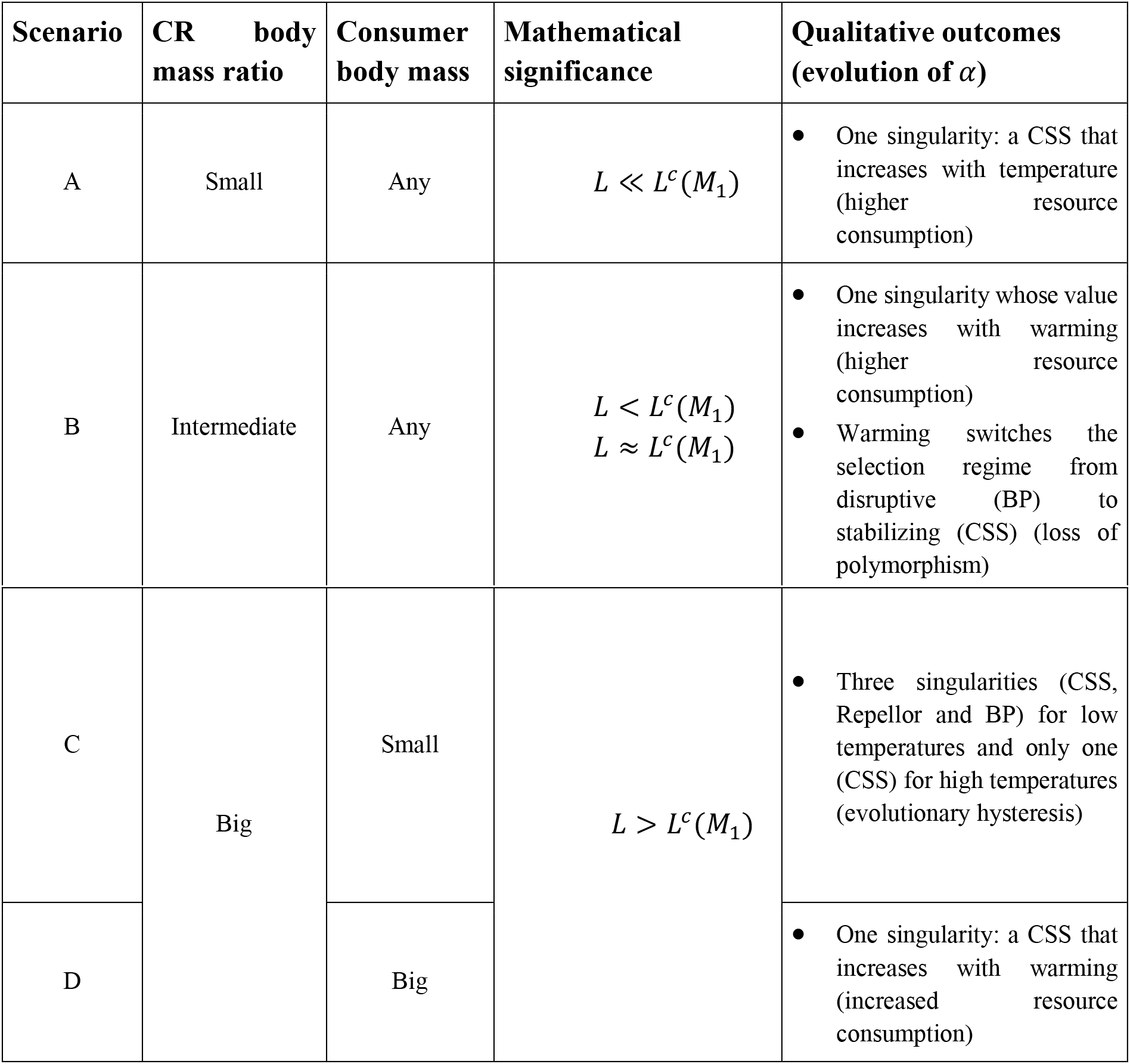
The four possible scenarios with their biological and mathematical significances. *L^c^*(*M*_1_) is the value of *L*, determined numerically and *M*_1_-dependent, around which important features of equation (H) change, leading to different evolutionary dynamics.

###### d. Nature of the evolutionary singularities

Let 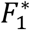 be the evolutionary singularity.

Non-invasibility corresponds mathematically to:

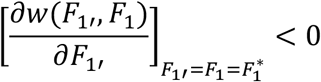

Moreover, we have the following expression obtained by derivation:

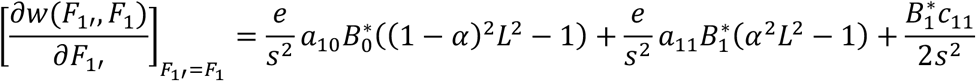

We used this expression to determine numerically trough Python software the invasibility properties.

Convergence corresponds mathematically to:

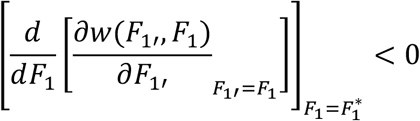

In the case of one singularity 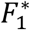, it can be shown without calculation that the singularity will be convergent stable.

###### Proof

Let *F* < *M*_0_ and 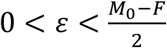, we have *w*(*F + ϵ, F*) > 0 because the mutant with trait (*F + ϵ*) has better attack rates on both the resource and the resident *F* and experiences less competition with the resident *F* than experienced by the resident (*C*_11_ = *c*_0_).

Hence, 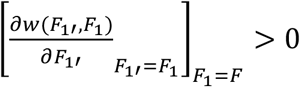.

In the same way, we have: 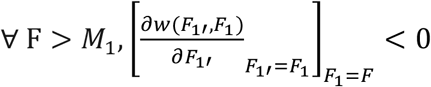

Moreover, the function:

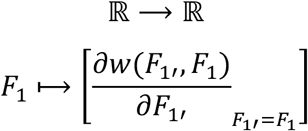

is continuous and only vanishes at 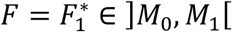.

Hence, 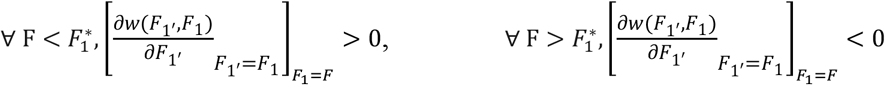 and 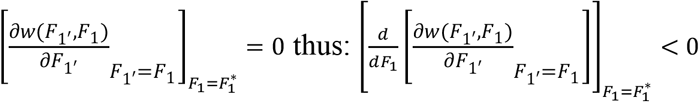

In the case of three singularities 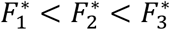, with the same kind of arguments, we know at least one singularity is convergent stable. It appears numerically that 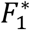 and 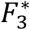 are convergent stable while 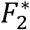 is not. This analysis gives the qualitative outcomes of warming for each scenario presented in **Table S1**.

###### e. Robustness check: variation of the conversion efficiency

The conversion efficiency is likely to vary with the feeding mode, with overall smaller values for herbivores and larger values for carnivores, as estimated by (Yodzis & Innes, 1992). We propose here to investigate the effect of warming on the consumer’s evolutionary dynamics (trait a) for a value of conversion efficiency that corresponds to herbivory (*e* = 0.45).

###### Value of *T_lim_*

The critical temperature value above which the consumer population cannot survive anymore increases with the conversion efficiency as indicated by equation (B).

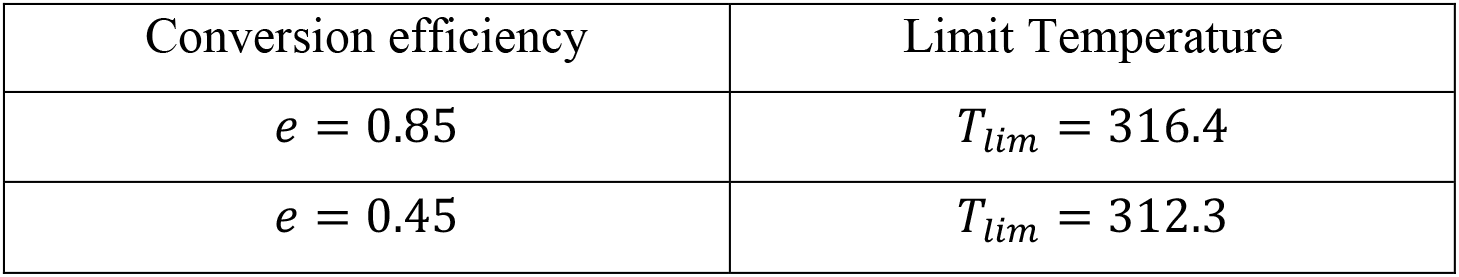

###### Evolutionary dynamics

Fig. B (akin Fig. 3) shows the evolutionary dynamics of the trait a according to temperature for the four scenarios (Table 1).

**Figure C:**
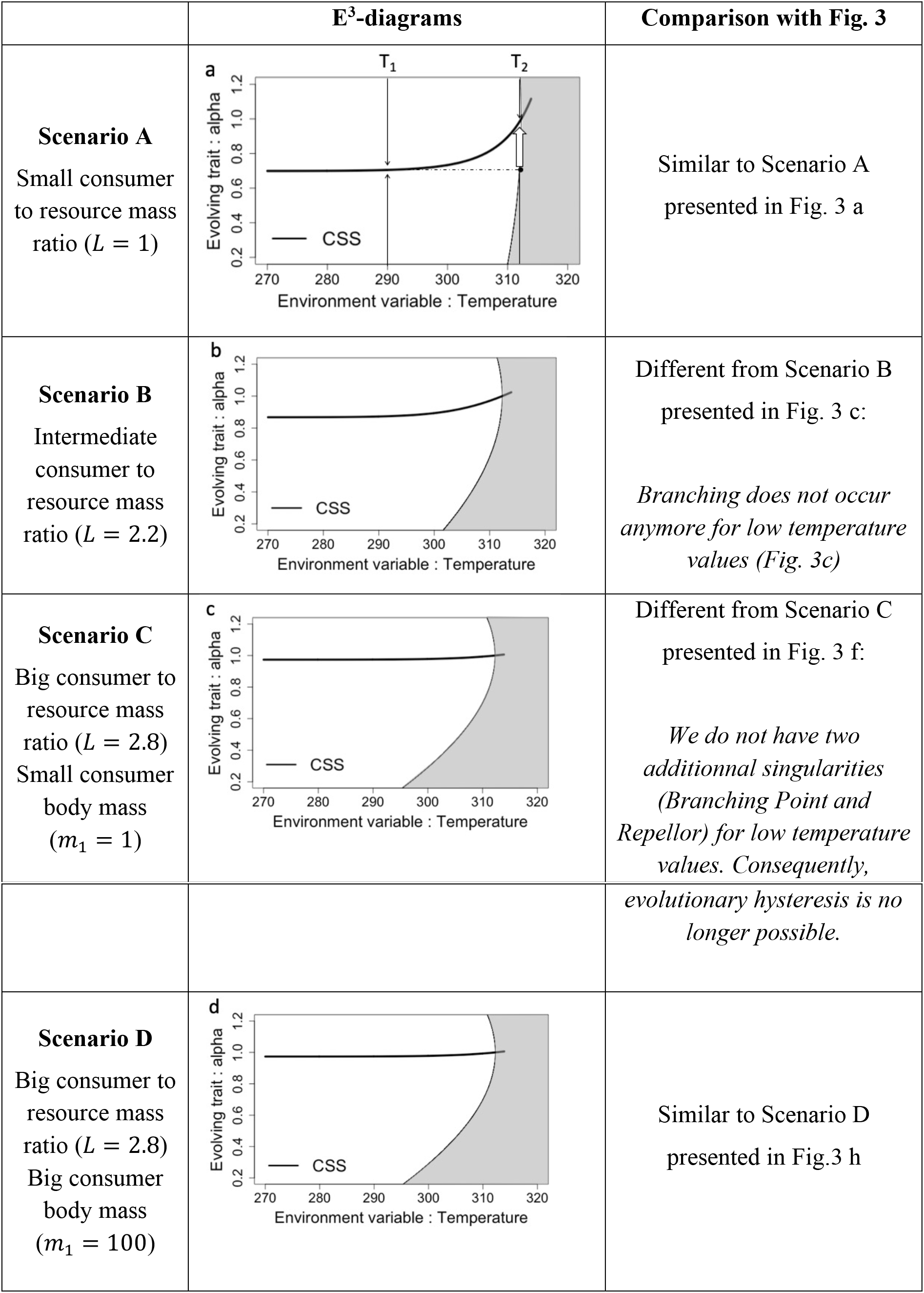
E^3^ diagrams corresponding respectively to scenarios A, B, C and D (**Table S1**). Grey areas are nonviability regions; the curves indicate the evolutionary singularities with the type of lines indicating their nature. **a.** Thin black arrows indicate the direction of evolution and the big white arrow indicates potential evolutionary rescue. Note that there is potential for evolutionary rescue in all scenarios although it is illustrated only in **a.**

This analysis reveals that some of the results observed for a higher value of conversion efficiency hold while others do not.

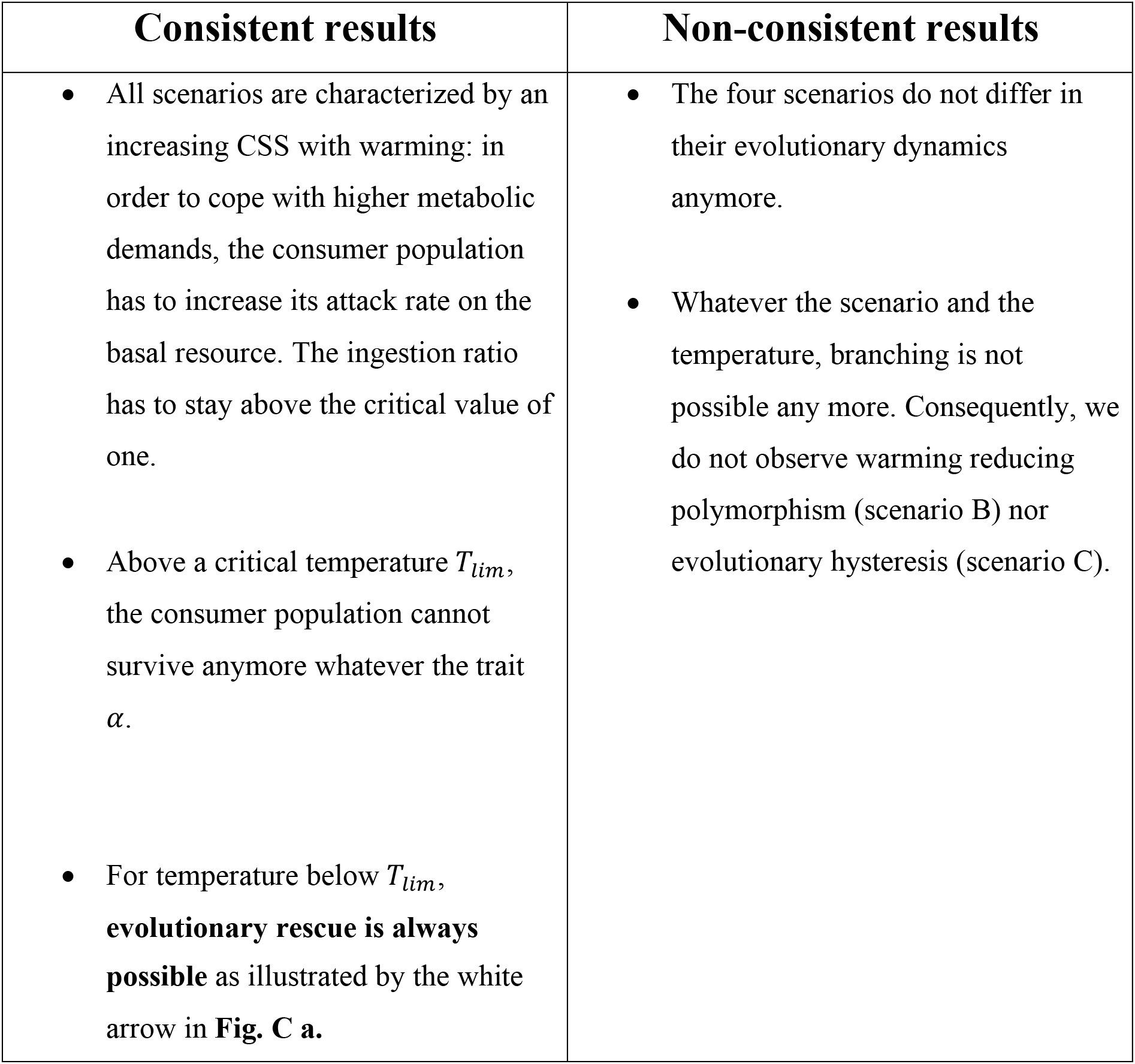

All in all, this robustness check confirms that evolutionary rescue is a potential mechanism by which evolution can impede the consumer population extinction given the temperature stays below a critical value *T_lim_*. However, some of our results, such as the potential for warming to dampen diversification processes (Scenario B, Fig. 3) seem to only apply for higher value of conversion efficiency. This can be interpreted as additional evidence that higher trophic levels are more vulnerable to warming (e.g. Binzer et al., 2012).

##### 4. The co-evolution of body mass and feeding preference

The separation between body mass and feeding preference evolution implemented so far may only be justified if one trait evolves much faster than the other (e.g. asymmetries in heritabilities). In other conditions, the two traits coevolve. We simulated two co-evolutionary scenarios that differ in mutation rates (10^−2^ for both traits in scenario 1 and 10^−2^ (resp. 10^−3^) for body mass (resp. feeding preference) in scenario 2). Warming goes from 280 to 316.4K (from 7 to 43°C, slightly below *T_lim_*). Evolutionary trajectories are shown on **Fig. S.** Evolutionary rescue occurs and enables food chain persistence. Note two important results. First, the two scenarios yield different evolutionary trajectories. The relative speed of evolution between the two evolving traits therefore affect evolutionary dynamics, possibly constraining evolutionary rescue. Second, while warming does not affect body mass when only body mass is allowed to evolve (equation (E), **Fig. 3**), it does under co-evolution. Here, warming exerts a selective pressure on the consumer feeding preference (see equation E) whose evolution, in turn, exerts a selective pressure on the consumer body mass. Consequently, both body mass and feeding preference are affected by warming (squares and triangles trajectories in **Fig. S1a**). Evolutionary rescue trajectories (**Fig. S1b**) then involve the effective evolution of both traits.

**Figure S2:**
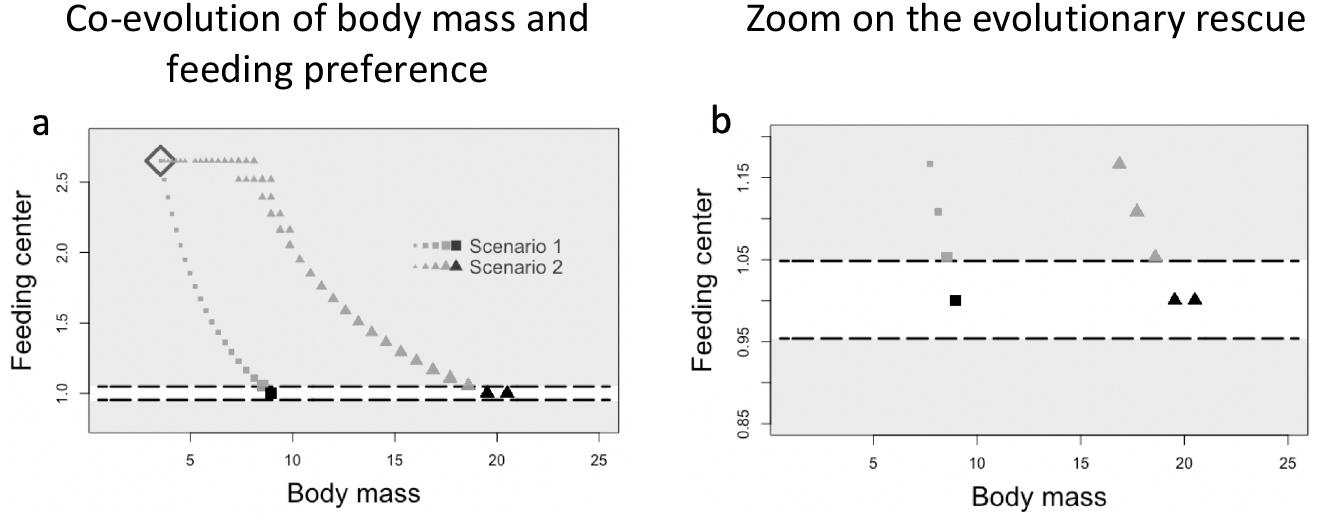
**a.** Co-evolution of body mass and feeding preference under two scenarios. **Scenario 1**: body mass and feeding preference evolve at rate ***μ* = 10^−2^** (squares). **Scenario 2**: body mass evolves at ***μ* = 10^−2^** while feeding preference evolves at ***μ* = 10^−3^** (triangles). Unfilled grey diamond: initial conditions. The simulations started at 280 K and the increase in symbol size indicates the direction of time. The biggest grey symbols thus correspond to the long-term selected phenotypes at 280 K. Once such a stable evolutionary state is reached, a warming to 316.4 K (almost maximal sustainable temperature) occurs. The consumer body mass and feeding preference, which were within the non-viability area at 316.4 K (light grey), evolve in response to warming. The newly selected phenotypes (black symbols) are within the viability area at 316.4 K (white), which indicates an evolutionary rescue process. **b.** Zoom (y-axis) to see the evolutionary rescue consecutive to warming. Note that the selected phenotypes at 280 K were one mutation away from being viable at 316.4 K. Parameter values are in **table 1**.

This result indicates that the eco-evolutionary dynamics within the complex multi-trophic network, where both traits coevolve, is likely to exhibit complex patterns not captured within the CR module. This complexity arises from the indirect interactions at play. Indirect interactions correspond for instance to interactions between the two evolving traits, as illustrated in this section. The many indirect ecological interactions occurring at the network scales also explain this additional complexity.

## Appendix S2: Complex multitrophic network model

Our community evolution model starts with a consumer feeding on the basal resource. The evolution of the consumer body mass and feeding preference follows a mutation/selection process leading to the emergence of a complex multi-trophic network of approximately 30 to 40 morphs and 4 to 5 trophic levels. A morph corresponds to an adaptive phenotype, that is to say, a couple (*body mass, feeling preference*).

### A. Definition of species diversity

At first, we only have one straightforward diversity measure: the trait diversity (i.e. the number of morphs in the trophic network at a given time). Because our model ignores genetic details and focuses on phenotypes, the definition of species is notoriously tricky. For lack of better criteria, we define species as clusters in the phenotypic space. The silhouette method (Rousseeuw, 1987) is used to determine the best number of clusters on the k-means clustering algorithm applied on the set of body masses and feeding preferences (*m_i_,/f_i_*) corresponding to the trait diversity. The number of clusters gives what we define as species diversity.

We consider a set of observations and a clustering. For each observation *i*, we note *C_i_* the cluster to which *i* is affected. For each observation *i*, the silhouette width *S*(*i*) is a measure of how much the observation is close to its cluster *C_i_* in comparison with other clusters. It is defined as follow:

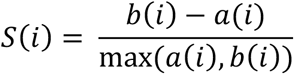

Where:

- *a*(*i*) is the average dissimilarity between *i* and the other observations in *C_i_*
- *b*(*i*) is the average dissimilarity between *i* and the observations in the closest cluster to *i, C*, with 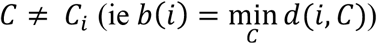

*S*(*i*) is between −1 and 1 and the more it is close to 1, the better is the affectation of in the clustering. We use 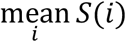 as a measure of the performance of a given clustering. We compare the different clustering obtained by the k-means algorithm for different values of k.

We implemented the algorithm in R-software using the packages “Cluster” by Martin Maechler, Peter Rousseeuw Anja Struyf, Mia Hubert, Kurt Hornik, Pierre Roudier, Juan Gonzalez (2017).

### B. Simulations

#### Random draw of mutant’s traits

Proportionally to the population densities distribution, a parent morph is chosen randomly at each mutation event. Mutant traits are then drawn from log-normal distributions centered on the parent’s traits. More precisely, mutantj’s traits *log*_10_(*m_j_*) (resp. *log*_10_(*f_j_*)) are randomly selected from a normal distribution of mean *log*_10_(*m_i_*) (resp. *log*_10_(*f_i_*)) and variance *σ*^2^. The value of *σ* (0.25) allows occasional big mutational steps: 5% of the mutations result in 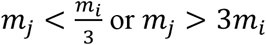.

#### Transition criterion

According to the scenario, each simulation follows a sequence of events.

- Scenarios NE: (1) the network is built up with a mutation rate *μ*; (2) evolution stops (*μ* = 0); (3) warming occurs; (4) simulation stops.
- Scenarios E: (1) the network is built up with a mutation rate *μ*; (2) warming occurs; (3) simulation stops.

Each transition is triggered once the transient dynamics are over. We consider these transient dynamics to be over when the ratio between the trait diversity standard deviation and mean (coefficient of variation) over a time window of 2.10^6^ mutation events is below 0.045.

This translates into:

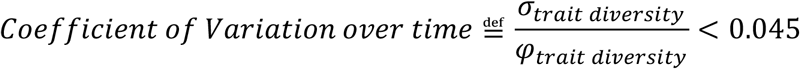

Where *σ_trait diversity_* is the standard deviation of the trait diversity over a time window of 2.10^6^ mutation events and *φ_trait diversity_* is the trait diversity mean over the same time period. Such a criterion allows for at least 95% of the observed trait diversity over this time window to be within the range of ±10& of the observed trait diversity mean over the same period.

#### Diversity before and after warming (persistence)

Due to the constant mutation/extinction events, diversity is subject to stochastic variation while we would like to have exactly one measure of diversity before and after warming to assess performance. Therefore, we take the diversity before (resp. after) as the average diversity over the time window of 2.10^6^ mutation events that satisfied the transition criterion before (resp. after) warming (*φ_trait diversity_*). Each simulation gives a diversity maintenance measure we call “persistence”, either calculated for trait or species diversity.

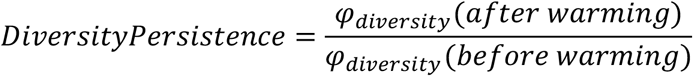

### C. Statistical analysis

We want to compare the diversity response to warming within our multi-trophic network with and without evolution.

For trait and species diversity, three models were fitted (*μ* = 10^−^, 10^−2^, 10^−3^) in order to contrast diversity persistence for different evolutionary scenarios (scenario E *μ* against scenario NE *μ*). 6 ANCOVAs were fitted with evolution as factor (two levels: “evolution” versus “no evolution”) and intensity of warming as quantitative variable (0,2,4, 6,8,10,20H). The model is written as follows:

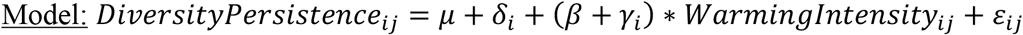

As indicated in **table S2**, index 1 (resp. index 2) corresponds to the level “no evolution” (resp. “evolution”). The level “no evolution” serves as a reference (*δ*_1_ = 0, *γ*_1_, = 0). Therefore, *δ*_2_ corresponds to the direct effect of evolution, while *γ*_2_ corresponds to the interaction term between evolution and warming or indirect effect of evolution. *β* describes the direct effect of warming.

**Table S2:**
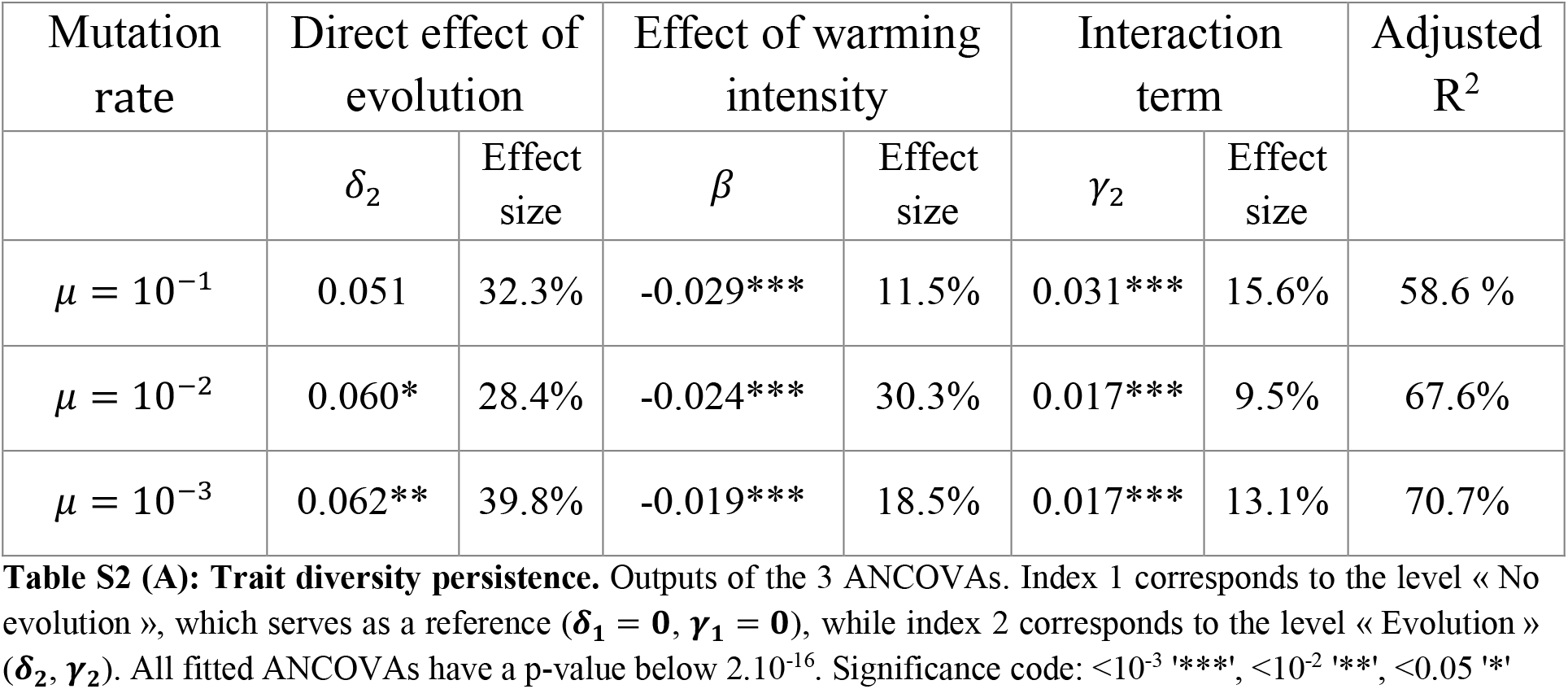
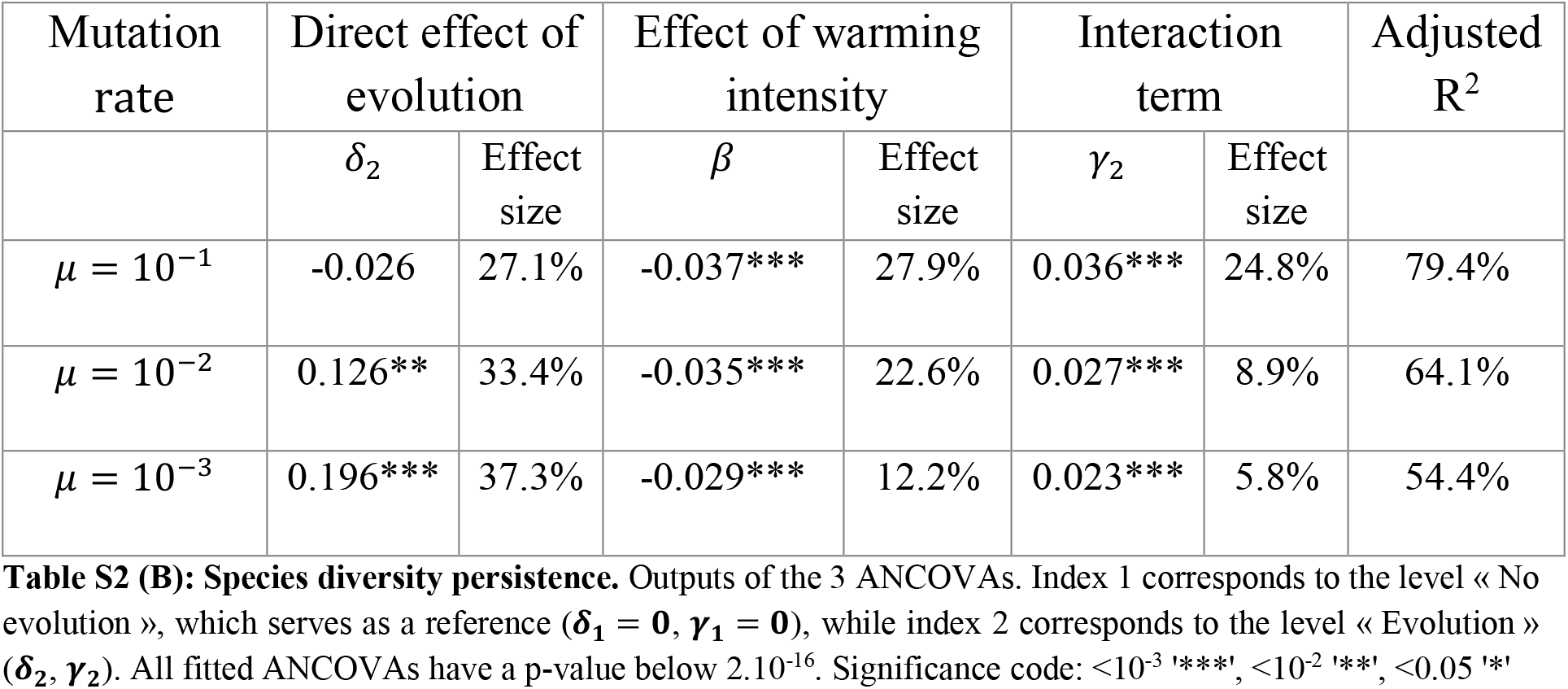
Diversity persistence in the complex multitrophic networks: statistical analysis summary for trait diversity (A) and species diversity (B)

An additional ANCOVA was fitted in order to compare the diversity persistence according to the mutation rate *μ*. Here, diversity persistence is explained by the mutation rate (*μ* = 10^−^’, 10^−2^,10^−3^) as a three-level factor and the intensity of warming (0, 2, 4, 6, 8, 10, 20 K) as a quantitative variable. The model is written as follows (but results are not shown because not significant):

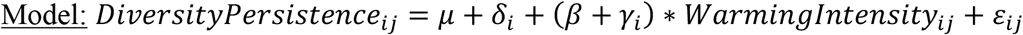

A graphical visualization showed all hypothesis for ANCOVA were verified for each fitted model. All statistical analyses were conducted using R-software (version 3.5.2). The “Anova” function of the R-package “car” was also used.

## Appendix S3: Simulation code

The C code given as supporting information (file: SimulationCode.c) corresponds to the code simulating complex multi-trophic networks for scenarios NE (i.e. without evolution). In order for the code to work, the folder where it is launched has to contain a folder with a correct path (see variable “path” L75 & input data L 149). In addition to this variable “path”, 3 other inputs are necessary (see L 149-152 of the code): a seed, a new temperature and a deltaTm (i.e. ^1^/*μ, μ* being the mutation rate).

In order to simulate scenarios E (with evolution), the following 2 changes are necessary:

(1): L 301: **while** (t<tend && S<Smax && S>0 && state<3)

Has to be replaced by:

**while** (t<tend && S<Smax && S>0 && state<2)

(2) L341-344:

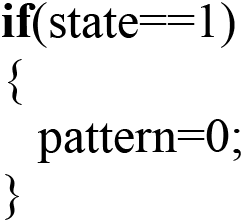

Has to be replaced by:

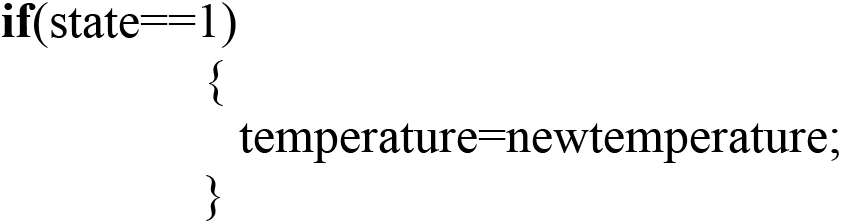

## References

Aljetlawi, A.A., Sparrevik, E., & Leonardsson, K. (2004). Prey-predator size-dependent functional response: Derivation and rescaling to the real world. Journal of Animal Ecology (Vol. 73). doi:/10.1111/j.0021-8790.2004.00800.x

Allhoff, K. T., Ritterskamp, D., Rall, B. C., Drossel, B., & Guill, C. (2015). Evolutionary food web model based on body masses gives realistic networks with permanent species turnover. Scientific Reports, 5(May), 10955. doi:10.1038/srep10955

Alvarez, S. A., Gibbs, S. J., Bown, P. R., Kim, H., Sheward, R. M., & Ridgwell, A. (2019, October 25). Diversity decoupled from ecosystem function and resilience during mass extinction recovery. Nature. Nature Publishing Group. doi:10.1038/s41586-019-1590-8

Barnosky, A. D., Matzke, N., Tomiya, S., Wogan, G. O. U., Swartz, B., Quental, T. B., … Ferrer, E. A. (2011). Has the Earth’s sixth mass extinction already arrived? Nature, 471(7336), 51–57. doi:10.1038/nature09678

Binzer, A., Guill, C., Brose, U., & Rall, B. C. (2012). The dynamics of food chains under climate change and nutrient enrichment. Philosophical Transactions of the Royal Society B: Biological Sciences, 367(1605), 2935–2944. doi:10.1098/rstb.2012.0230

Brose, U., Ehnes, R. B., Rall, B. C., Vucic-Pestic, O., Berlow, E. L., & Scheu, S. (2008). Foraging theory predicts predator-prey energy fluxes. Journal of Animal Ecology, 77(5), 1072–1078. doi:10.1111/j.1365-2656.2008.01408.x

Brown, J. H., Gillooly, J. F., Allen, A. P., Savage, V. M., & West, G. B. (2004). Toward a metabolic theory of ecology. Ecology, 85(7), 1771–1789. doi:Doi 10.1890/03-9000

Buckley, J., Butlin, R. K., & Bridle, J. R. (2012). Evidence for evolutionary change associated with the recent range expansion of the British butterfly, Aricia agestis, in response to climate change. Molecular Ecology, 21(2), 267–280. doi:10.1111/j.1365-294X.2011.05388.x

Carlson, S. M., Cunningham, C. J., & Westley, P. A. H. (2014). Evolutionary rescue in a changing world. Trends in Ecology and Evolution, 29(9), 521–530. doi:10.1016/j.tree.2014.06.005

Carroll, S. P., Loye, J. E., Dingle, H., Mathieson, M., Famula, T. R., & Zalucki, M. P. (2005). And the beak shall inherit - Evolution in response to invasion. Ecology Letters, 8(9), 944–951. doi:10.1111/j.1461-0248.2005.00800.x

Colwell, R. K., & Rangel, T. F. (2009). Hutchinson’s duality: The once and future niche. Proceedings of the National Academy of Sciences, 106(Supplement_2), 19651–19658. doi:10.1073/pnas.0901650106

Dakos, V., Matthews, B., Hendry, A. P., Levine, J., Loeuille, N., Norberg, J., … De Meester, L. (2019). Ecosystem tipping points in an evolving world. Nature Ecology and Evolution, 5(3), 355–362. doi:10.1038/s41559-019-0797-2

Daufresne, M., Lengfellner, K., & Sommer, U. (2009). Global warming benefits the small in aquatic ecosystems. Proceedings of the National Academy of Sciences of the United States of America, 106(31), 12788–12793. doi:10.1073/pnas.0902080106

Dieckmann, U., & Law, R. (1996). The dynamical theory of coevolution: a derivation from stochastic ecological processes. Journal of Mathematical Biology, 54(5-6), 579–612. doi:10.1007/BF02409751

Dieckmann, U., Marrow, P., & Law, R. (1995). Evolutionary cycling in predator-prey interactions: Population dynamics and the red queen. Journal of Theoretical Biology, 176(1), 91–102. doi:10.1006/jtbi.1995.0179

Ellner, S. P., & Becks, L. (2011). Rapid prey evolution and the dynamics of two-predator food webs. Theoretical Ecology, 4(2), 133–152. doi:10.1007/s12080-010-0096-7

Englund, G., Öhlund, G., Hein, C. L., & Diehl, S. (2011). Temperature dependence of the functional response. Ecology Letters, 14(9), 914–921. doi:10.1111/j.1461-0248.2011.01661.x

Eshel, I. (1983). Evolutionary and continuous stability. Journal of Theoretical Biology, 105(1), 99–111. doi:10.1016/0022-5193(83)90201-1

Ferriere, R., & Legendre, S. (2012). Eco-evolutionary feedbacks, adaptive dynamics and evolutionary rescue theory. Philosophical Transactions of the Royal Society B: Biological Sciences, 568(1610), 20120081–20120081. doi:10.1098/rstb.2012.0081

Fugère, V., Hébert, M. P., da Costa, N. B., Xu, C. C. Y., Barrett, R. D. H., Beisner, B. E., … Gonzalez, A. (2020). Community rescue in experimental phytoplankton communities facing severe herbicide pollution. Nature Ecology and Evolution, 4(4), 578–588. doi:10.1038/s41559-020-1134-5

Fussmann, K. E., Schwarzmüller, F., Brose, U., Jousset, A., & Rall, B. C. (2014). Ecological stability in response to warming. doi:10.1038/NCLIMATE2134

Geritz, S. A. H., Metz, J. A. J., Kisdi, É., & Meszéna, G. (1997). Dynamics of adaptation and evolutionary branching. Physical Review Letters, 78(10), 2024–2027. doi:10.1103/PhysRevLett.78.2024

Gomulkiewicz, R., & Holt, R. D. (1995a). When does Evolution by Natural Selection Prevent Extinction? Source: Evolution BRIEF COMMUNK]AlION S Evolution, 49(491), 201–207. Retrieved from http://www.jstor.org/stable/2410305

Gomulkiewicz, R., & Holt, R. D. (1995b). When does Evolution by Natural Selection Prevent Extinction? Evolution, 49(1), 201–207. doi:10.2307/2410305

Hendry, A. P., Farrugia, T. J., & Kinnison, M. T. (2008). Human influences on rates of phenotypic change in wild animal populations. Molecular Ecology, 17(1), 20–29. doi:10.1111/j.1365-294X.2007.03428.x

Koch, H., Frickel, J., Valiadi, M., & Becks, L. (2014). Why rapid, adaptive evolution matters for community dynamics. Frontiers in Ecology and Evolution, 2, 17. doi:10.3389/fevo.2014.00017

Lavergne, S., Mouquet, N., Thuiller, W., & Ronce, O. (2010). Biodiversity and Climate Change: Integrating Evolutionary and Ecological Responses of Species and Communities. Annual Review of Ecology, Evolution, and Systematics, 41(1), 321–350. doi:10.1146/annurev-ecolsys-102209-144628

Lindeman, R. (1942). The Trophic-Dynamic aspect of ecology. Ecology.

Loeuille, N., & Loreau, M. (2005). Evolutionary emergence of size-structured food webs. Proceedings of the National Academy of Sciences, 102(16), 5761–5766. doi:10.1073/pnas.0408424102

Loeuille, Nicolas. (2019). Eco-evolutionary dynamics in a disturbed world: implications for the maintenance of ecological networks. F1000Research, 8(0), 97. doi:10.12688/f1000research.15629.1

Macarthur, R., & Levins, R. (1967). The Limiting Similarity, Convergence, and Divergence of Coexisting Species. The American Naturalist, 101(921), 377–385. doi:10.1086/282505

Metz, J. A. J., Nisbet, R. M., & Geritz, S. A. H. (1992). How should we define “fitness” for general ecological scenarios? Trends in Ecology and Evolution (Vol. 7). doi:/10.1016/0169-5347(92)90073-K

Naisbit, R. E., Kehrli, P., Rohr, R. P., & Bersier, L. F. (2011). Phylogenetic signal in predator–prey body-size relationships. Ecology, 92(12), 2183–2189. doi:10.1890/10-2234.1

Norberg, J., Urban, M. C., Vellend, M., Klausmeier, C. A., & Loeuille, N. (2012). Eco-evolutionary responses of biodiversity to climate change. Nature Climate Change, 2(10), 747–751. doi:10.1038/nclimate1588

O’Gorman, E. J., Zhao, L., Pichler, D. E., Adams, G., Friberg, N., Rall, B. C., … Woodward, G. (2017). Unexpected changes in community size structure in a natural warming experiment. Nature Climate Change, 7(9), 659–663. doi:10.1038/nclimate3368

Olsen, E. M., Heino, M., Lilly, G. R., Morgan, M. J., Brattey, J., Ernande, B., & Dieckmann, U. (2004). Maturation trends indicative of rapid evolution preceded the collapse of northern cod. Nature, 428(6986), 932–935. doi:10.1038/nature02430

Osmond, M. M., Otto, S. P., & Klausmeier, C. A. (2017). When Predators Help Prey Adapt and Persist in a Changing Environment. The American Naturalist, 190(1), 83–98. doi:10.1086/691778

Parmesan, Camilla. (2006). Ecological and Evolutionary Responses to Recent Climate Change. Annual of Ecology, Evolution and Systematics, 37, 637–669. doi:10.2307/annurev.ecolsys.37.091305.30000024

Parmesan, Camille, & Yohe, G. (2003). A globally coherent fingerprint of climate change impacts across natural systems. Nature, 421(6918), 37–42. doi:10.1038/nature01286

Pateman, R. M., Hill, J. K., Roy, D. B., Fox, R., & Thomas, C. D. (2012). Temperature-Dependent Alterations in Host Use Drive Rapid Range Expansion in a Butterfly. Science, 336(6084), 1028–1030. doi:10.1126/science.1216980

Pearson, R. G., Dawson, T. P., Berry, P. M., & Harrison, P. A. (2002). SPECIES: A spatial evaluation of climate impact on the envelope of species. Ecological Modelling, 154(3), 289–300. doi:10.1016/S0304-3800(02)00056-X

Petchey, O. L., McPhearson, P. T., Casey, T. M., & Morin, P. J. (1999). Environmental warming alters food-web structure and ecosystem function. Nature, 402(6757), 69–72. doi:10.1038/47023

Peters, R. H. (1983). The ecological implications of body size. Cambridge University Press. Cambridge University Press.

Pounds, J. A., Fogden, M. P. L., & Campbell, J. H. (1999). Biological response to climate change on a tropical mountain. Nature (Vol. 398). doi:/10.1038/19297

Rall, B. C., Brose, U., Hartvig, M., Kalinkat, G., Schwarzmüller, F., Vucic-Pestic, O., & Petchey, O. L. (2012). Universal temperature and body-mass scaling of feeding rates. Philosophical Transactions of the Royal Society B: Biological Sciences, 367(1605), 2923–2934. doi:10.1098/rstb.2012.0242

Rall, B. Ö. C., Vucic-Pestic, O., Ehnes, R. B., EmmersoN, M., & Brose, U. (2010). Temperature, predator-prey interaction strength and population stability. Global Change Biology, 16(8), 2145–2157. doi:10.1111/j.1365-2486.2009.02124.x

Sheridan, J. A., & Bickford, D. (2011). Shrinking body size as an ecological response to climate change. Nature Climate Change. doi:10.1038/nclimate1259

Singer, M. C., & Parmesan, C. (2018). Lethal trap created by adaptive evolutionary response to an exotic resource. Nature, 557(7704), 238–241. doi:10.1038/s41586-018-0074-6

Smith, T. B., Bruford, M. W., & Wayne, R. K. (1993). The Preservation of Process: The Missing Element of Conservation Programs The preservation of process: the missing element conservation programs. Source: Biodiversity Letters Biodiversity Letters, 1(1). Retrieved from http://www.jstor.org/stable/2999740

Stockwell, C. A., Hendry, A. P., & Kinnison, M. T. (2003). Contemporary evolution meets conservation biology. Trends in Ecology and Evolution, 18(2), 94–101. doi:10.1016/S0169-5347(02)00044-7

Suding, K. N., & Hobbs, R. J. (2009). Threshold models in restoration and conservation: a developing framework. Trends in Ecology and Evolution, 24(5), 271–279. doi:10.1016/j.tree.2008.11.012

Tylianakis, J. M., Didham, R. K., Bascompte, J., & Wardle, D. A. (2008). Global change and species interactions in terrestrial ecosystems. Ecology Letters. doi:10.1111/j.1461-0248.2008.01250.x

Uszko, W., Diehl, S., Englund, G., & Amarasekare, P. (2017). Effects of warming on predator-prey interactions - a resource-based approach and a theoretical synthesis. Ecology Letters. doi:10.1111/ele.12755

Visser, M. E., & Gienapp, P. (2019, June 22). Evolutionary and demographic consequences of phenological mismatches. Nature Ecology and Evolution. Nature Publishing Group. doi:10.1038/s41559-019-0880-8

Vucic-Pestic, O., Rall, B. C., Kalinkat, G., & Brose, U. (2010). Allometric functional response model: Body masses constrain interaction strengths. Journal of Animal Ecology, 79(1), 249–256. doi:10.1111/j.1365-2656.2009.01622.x

Weinbach, A., Allhoff, K., Thebault, E., Massol, F., & Loeuille, N. (2017). Selective effects of temperature on body mass depend on trophic interactions and network position. BioRxiv, 233742. doi:10.1101/233742

Yachi, S., & Loreau, M. (1999). Biodiversity and ecosystem productivity in a fluctuating environment: the insurance hypothesis. Proceedings of the National Academy of Sciences of the United States of America, 96(4), 1463–8. doi:10.1073/PNAS.96.4.1463

Yamamichi, M., & Miner, B. E. (2015). Indirect evolutionary rescue: Prey adapts, predator avoids extinction. Evolutionary Applications, 8(8), 787–795. doi:10.1111/eva.12295

Yodzis, P., & Innes, S. (1992). Body Size and Consumer-Resource Dynamics. Source: The American Naturalist (Vol. 139). Retrieved from https://about.jstor.org/terms

## References

Yodzis, P., & Innes, S. (1992). Body Size and Consumer-Resource Dynamics. Source: The American Naturalist (Vol. 139).

## References

Rousseeuw, P. J. (1987). Silhouettes: A graphical aid to the interpretation and validation of cluster analysis. Journal of Computational and Applied Mathematics, 20, 53–65. doi:10.1016/0377-0427(87)90125-7

